# Personalized Network Modeling of the Pan-Cancer Patient and Cell Line Interactome

**DOI:** 10.1101/806596

**Authors:** Rupam Bhattacharyya, Min Jin Ha, Qingzhi Liu, Rehan Akbani, Han Liang, Veerabhadran Baladandayuthapani

## Abstract

**Purpose:** Personalized network inference on diverse clinical and *in vitro* model systems across cancer types can be used to delineate specific regulatory mechanisms, uncover drug targets and pathways, and develop individualized predictive models in cancer.

**Datasets and methods:** We developed TransPRECISE, a multi-scale Bayesian network modeling framework, to analyze the pan-cancer patient and cell line interactome to identify differential and conserved intra-pathway activities, globally assess cell lines as representative models for patients and develop drug sensitivity prediction models. We assessed pan-cancer pathway activities for a large cohort of patient samples (>7700) from The Cancer Proteome Atlas across ≥30 tumor types and a set of 640 cancer cell lines from the M.D. Anderson Cell Lines Project spanning16 lineages, and ≥250 cell lines’ response to >400 drugs.

**Results:** TransPRECISE captured differential and conserved proteomic network topologies and pathway circuitry between multiple patient and cell line lineages: ovarian and kidney cancers shared high levels of connectivity in the hormone receptor and receptor tyrosine kinase pathways, respectively, between the two model systems. Our tumor stratification approach found distinct clinical subtypes of the patients represented by different sets of cell lines: head and neck patient tumors were classified into two different subtypes that are represented by head and neck and esophagus cell lines, and had different prognostic patterns (456 vs. 654 days of median overall survival; P=0.02). The TransPRECISE-based sample-specific pathway scores achieved high predictive accuracy for drug sensitivities in cell lines across multiple drugs (median AUC >0.8).

**Conclusion:** Our study provides a generalizable analytical framework to assess the translational potential of preclinical model systems and guide pathway-based personalized medical decision-making, integrating genomic and molecular data across model systems.

## INTRODUCTION

Precision medicine aims to improve clinical outcomes by optimizing treatment to each individual patient. The rapid accumulation of large-scale pan-omic molecular data across multiple cancers on patients (ICGC,^1^ TCGA,^2^ PCAWG,^3^ TCPA^4,5^) and model systems (GDSC,^6^ CCLE,^7^ MCLP^8^), along with extensive drug profiling data (NCI60,^9^ LINCS,^10^ CMAP,^11–13^ DepMap^14^) have generated information-rich and diverse community resources with major implications for translational research in oncology.^15^ However, a major challenge remains: to bridge anticancer pharmacological data to large-scale omics in the paradigm wherein patient heterogeneity is leveraged and inferred through rigorous and integrative data-analytic approaches across patients and model systems.

Complex diseases, such as cancer, are often characterized by small effects in multiple genes and proteins that are interacting with each other by perturbing downstream cellular signaling pathways.^16–18^ It is well established that complex molecular networks and systems are formed by a large number of interactions of genes and their products operating in response to different cellular conditions and cell environment, i.e., model systems.^19^ To date, most, if not all approaches for mechanism and drug discovery have been constrained by the biological system^20,21^ (patients or cell-lines), specific cancer lineage,^22,23^ or by prior knowledge of specific genomic alterations.^24,25^ Hence, there is a critical need for robust computational methods that integrate molecular profiles across large cohorts of patients and model systems from multiple lineages in an unbiased data-driven manner to delineate specific regulatory mechanisms, uncover drug targets and pathways, and develop individualized predictive models in cancer.

We have recently developed a network-based framework called PRECISE (personalized cancer-specific integrated network estimation model) to estimate cancer-specific networks, infer patient-specific networks, and elicit interpretable pathway-level signatures.^26^ Using a large cohort of patients (>7700) from TCGA across 30+ tumor types, we have shown that PRECISE identifies pan-cancer commonalities and differences in proteomic network biology within and across tumors, allows robust tumor stratification that is both biologically and clinically informative, and has superior prognostic power compared to multiple existing approaches.^26^ In this paper, we present translational PRECISE (TransPRECISE, in short), a generalization of the PRECISE framework, to establish the translational relevance of these pathway signatures. Briefly, TransPRECISE uses a multi-scale Bayesian modeling strategy that infers *de novo* differential and conserved networks of intra-pathway circuitry between the two biological systems (patients and cell lines) for multiple cancers. Further, it identifies cell line “avatars” for patients based on pathway activities, and also develops machine learning based predictive models for drug sensitivity in both cell lines and patients to potentially guide pathway-based individualized medical decision-making. We also have developed an online, publicly available, comprehensive interactive database and visualization tool of our findings along with software code (https://rupamb.shinyapps.io/transprecise).

## DATASETS AND METHODS

### Cancer patients’ proteomic data

We used a dataset of 7714 patient samples across 31 different cancer types available from the Cancer Proteome Atlas (TCPA)^4,5^ (Supplementary Table (ST) S1). TCPA offers reverse-phase protein array (RPPA)-based proteomics datasets, profiled using extensively validated antibodies to nearly 200 proteins and phosphoproteins. The functional space of the antibodies covers major functional and signaling pathways relevant to human cancers. For this work, we used a total of 12 pathways, including DNA damage response, EMT, hormone signaling, apoptosis, TSC/mTOR, and RAS/MAPK (ST S2).

### Cancer cell lines’ proteomic and drug sensitivity data

We used RPPA-based protein expression data for cell lines available via the MD Anderson Cell Lines Project (MCLP).^8^ In set of 640 cancer cell lines spanning across 16 lineages, each cell line has RPPA expression data based on the same set of proteins as in the patient tumors (ST S3). Additionally, we used drug sensitivity data from the Genomics of Drug Sensitivity in Cancer (GDSC)^6^ database, with the sensitivity of 481 drugs assessed on a subset of 254 cell lines (ST S4). For the entirety of this paper, we denote cell line samples in lowercase and patient samples in uppercase letters.

### TransPRECISE framework

The TransPRECISE implementation can broadly be classified into three modules (Figure 1). The first module takes as input the combined proteomics data from patients and cell lines (as described above). The second module implements the PRECISE modeling framework that consists of three steps. In Step 1, for a specific cancer type, TransPRECISE uses a Bayesian graphical regression model to estimate the population-level network structure among proteins belonging to a particular pathway for a specific cancer type. In Step 2, these population level networks are then de-convolved to obtain sample-specific networks by evaluating the extremities of sample-specific protein expression densities; where-in nodes of the cancer-specific networks are labeled by patient-specific protein activity statuses evaluated under the cancer-specific network topologies. Finally, the sample-specific interaction structure of the proteins is further summarized to obtain TransPRECISE scores that quantify pathway activities corresponding to the pathway’s neutral, activated or suppressed status in an individual sample (both patient tumors and cell-lines). The model specific parameterization and inferential strategies are described in Supplementary Materials (SM) Sections S1.2-S1.5.

**Figure 1.**
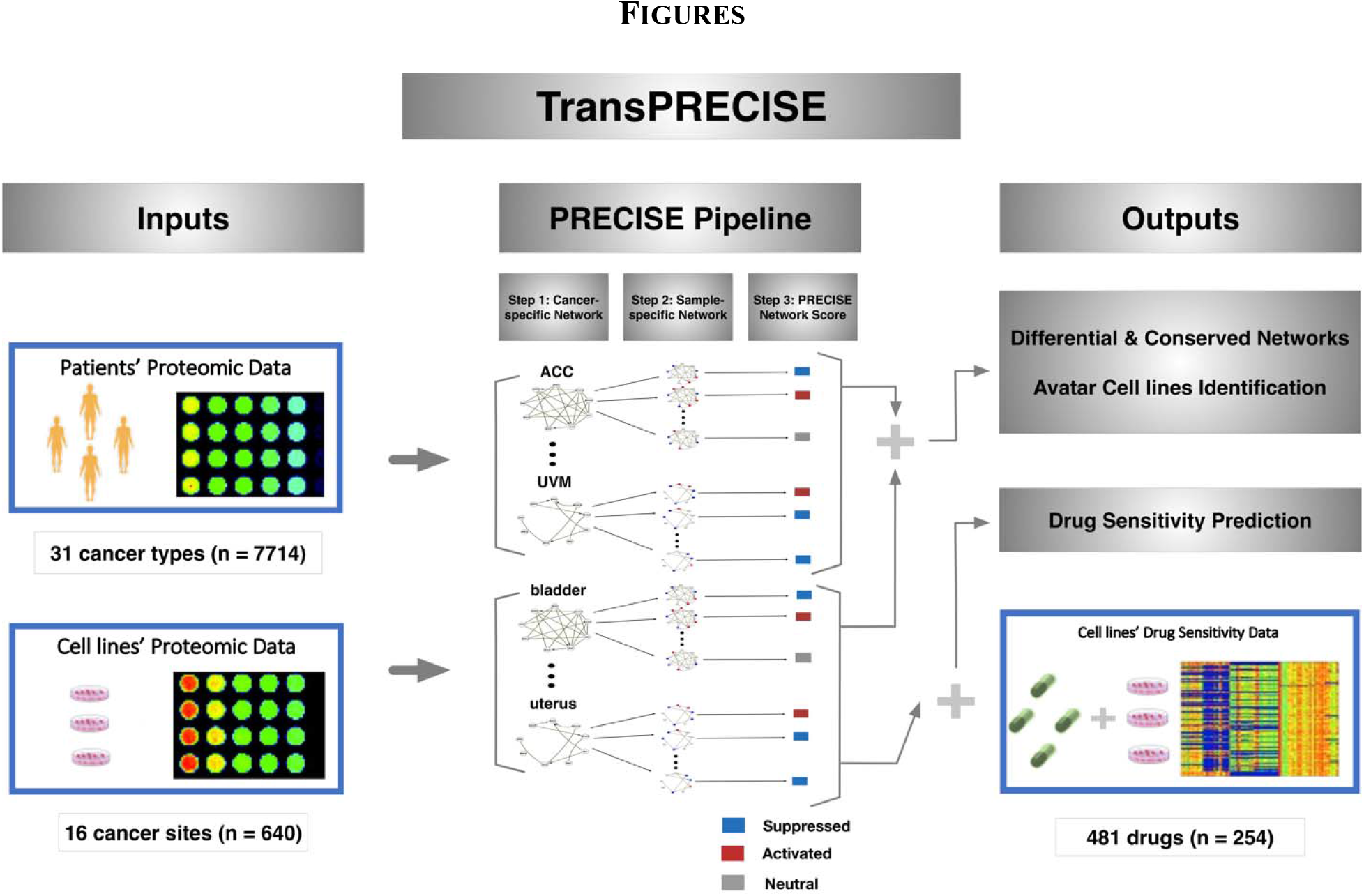
Overview of the TransPRECISE framework. The first step of TransPRECISE involves running the PRECISE pipeline on two sets of RPPA protein expression data—namely, cancer patients (7714 samples across 31 different cancer types) and cancer cell lines (640 samples across 16 different cancer tissues). For each combination of 47 cancer types across cell lines and patients and the 12 pathways, the PRECISE procedure is executed in three consecutive steps: fitting cancer-specific protein networks using Bayesian graphical regression, deconvolving these cancer-specific networks to fit sample-specific pathways networks, and aggregating the sample-specific networks to obtain calibrated TransPRECISE scores and pathway activity status. The obtained cancer-specific networks are then compared across patients and cell lines for each pathway towards a pan-cancer identification of differential and conserved pathway activities. The TransPRECISE scores are converted to sample-specific network aberration scores, and those are used to identify avatar cell lines for patient tumors: both at a global pathway level and at a pathway-specific interaction level. Finally, the network aberration scores are also used alongside drug sensitivity data for the cell lines for *in vivo* drug sensitivity prediction training, and the trained models are used on the patient network aberration scores for *in silico* drug sensitivity prediction.

## RESULTS

### Differential and conserved rewiring and circuitry of cancer-specific networks

Using the *de-novo* cancer-specific population-level networks (from Step 1 of TransPRECISE), we evaluated intra-pathway edge rewiring across lineages of the two model systems to identify highly conserved and differential edges, and to link patient and cell line tumor types by measuring intra-pathway circuitry.

#### Network rewiring across model systems

We determined the extent to which protein-protein edges in each of the pathways were shared across tumor sites in the patients and the cell lines. We found highly conserved edges across lineages for both cell lines and patients (Figure 2 and Supplementary Figures (SFs) S1-S10). All of the 12 pathways had at least one link that was shared across more than 20 lineages among the patient cancer types, and 11 pathways (with the exception of hormone signaling) had at least one link that was shared across more than 8 lineages among the cell line lineages. The conserved edges were further classified into three categories: patients-cell lines, (b) patients only, and (c) cell-lines only. For category (a), we identified significant regulation of CCNE2-FOXM1 (10 cell line lineages, 17 patient cancer types) in a cell cycle, CTNNB1-SERPINE1 (8 cell line lineages, 17 patient cancer types) in EMT, and RB1-RPS6 (8 cell line lineages, 20 patient cancer types) in TSC/mTOR pathways, respectively.

**Figure 2.**
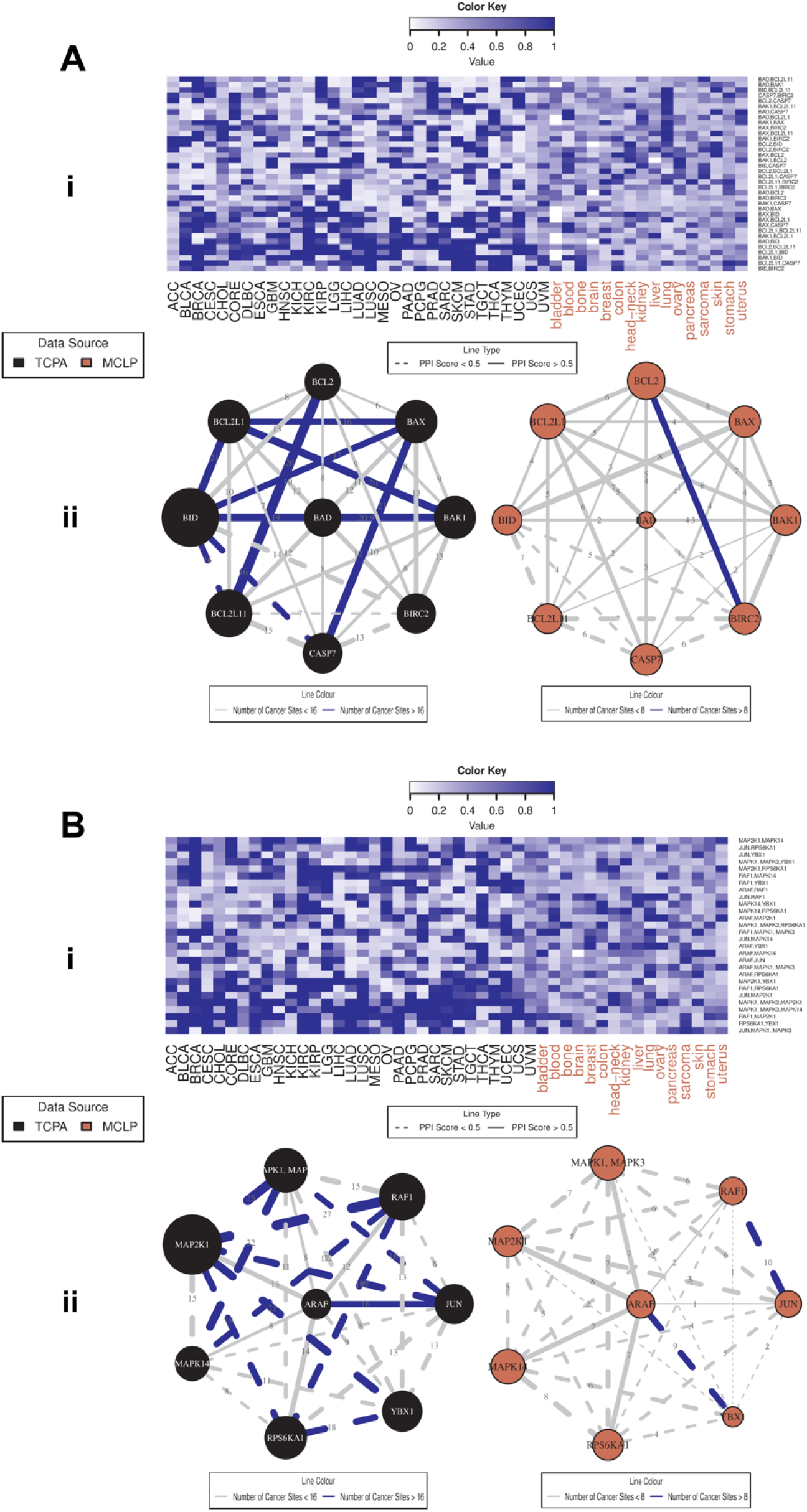
Pan-cancer summary of protein networks for apoptosis (A) and RAS/MAPK (B) pathways. i. Heatmap depicting strengths of all possible protein-protein edges within the pathway, across all 47 patient and cell line tumor lineages, quantified by the posterior inclusion probabilities of the edges based on the fitted Bayesian graphical regression model. ii. Left panel exhibits a network with its edges weighted and labeled by the edge consistencies (ECs), which are quantified by the number of patient tumor types holding that particular edge with a posterior probability >0.5, also presenting the *a priori* known strength of the edge using the protein-protein interaction score from the STRING database. The right panel is the corresponding network across cell line cancers.

#### Linking tumor types between model systems based on network circuitry

We investigated the shared cross-signaling between cell line and patient tumor types. As a measure of the level of cross-signaling of a specific pathway network, we defined the connectivity score (CS) as the ratio of the observed number of edges in a given network to the total number of possible edges in the pathway, as more edges imply a higher level of cross-signaling within a pathway (ST S6). In addition, we quantified the level of significance for the observed CS value by comparing it with CS values obtained from random permutation of the network, called randomCS; lower values of randomCS provide evidence against the observed CS value being obtained under random chance (SM Section S1.3). Based on the randomCS, we evaluated the similarity between cell line and patient tumor types in terms of network cross-signaling. Specifically, we declared two lineages were similar for a pathway if both of them showed high levels of cross-signaling (i.e., low randomCS proportions). Some key triplets of {cell line-pathways-patient} are summarized in Figure 3. Ovary and uterus cell lines were connected via the hormone signaling (breast) pathway with BRCA; lung, kidney, and stomach-oesophagus cell lines were linked together with two clusters of patient cancers (KICH, KIRP, PRAD, LGG and LUSC, UCEC, STAD) via the RTK pathway (Figure 3A). Further, there were some clear confirmations of conservation of activities across model systems within cancer tissues (Figure 3B), some specific examples being {bladder-core reactive-BLCA}, {kidney-RTK-KICH & KIRP}, {kidney-hormone receptor-KIRC}, {ovary-hormone signaling-OV}, and {stomach-hormone receptor-ESCA & STAD}.

**Figure 3.**
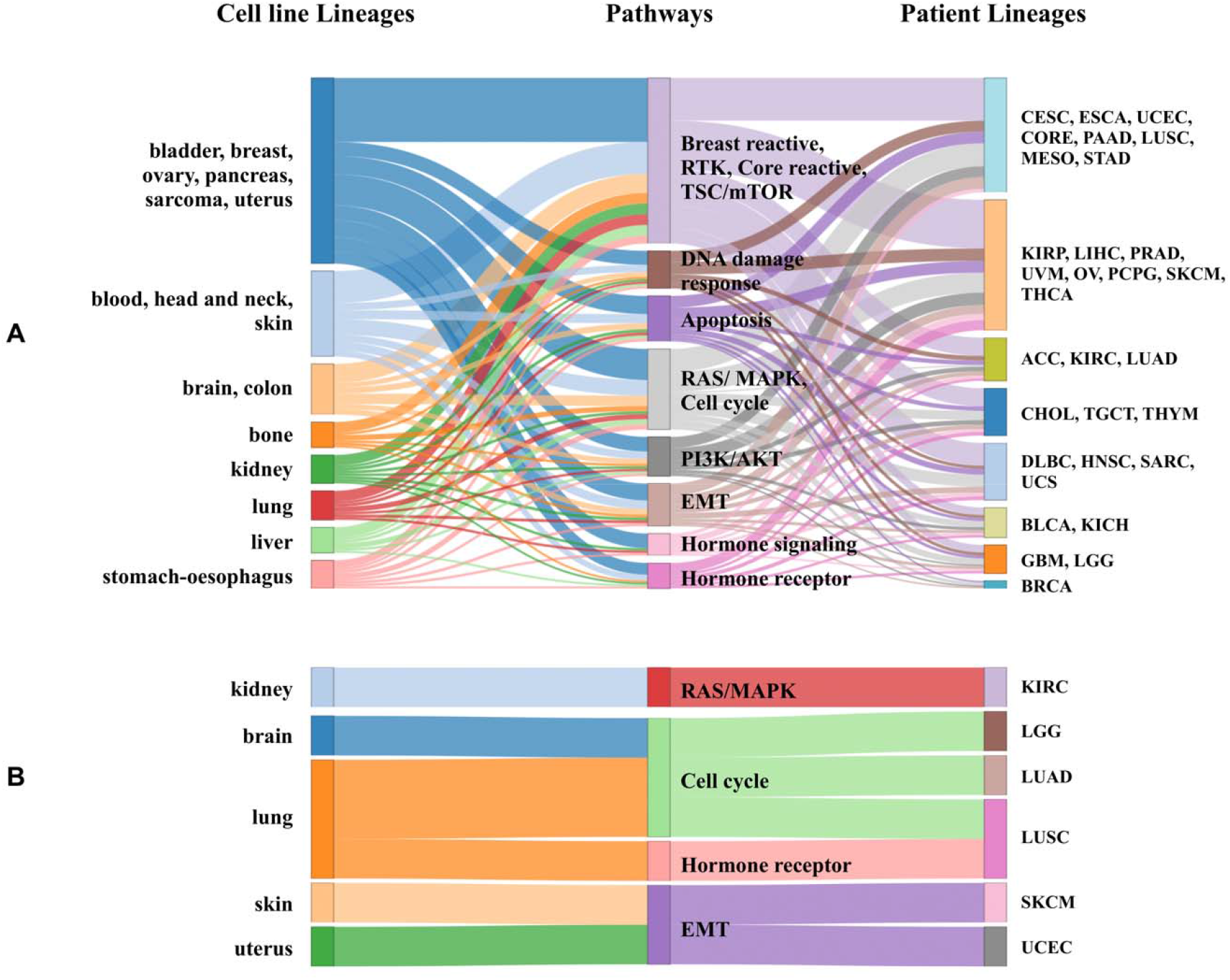
Sankey diagrams for patient and cell line cancers with conserved pathway-specific connectivity. A. The columns contain cell line cancers, pathways, and patient cancers from left to right, respectively. A cell line cancer tissue is connected to a pathway if the connectivity score (CS) for that cancer type-pathway pair (defined as the proportion of edges out of all possible undirected edges in the pathway that are held by that cancer type) is more than 900 out of 1000 randomCS values computed for that cancer type, with repeated random selection of the same number of proteins as in the pathway from the pool of all proteins across the 12 pathways. The connection between a patient cancer type to a pathway is also determined by the same rule. The length of the middle (pathway) column pieces indicate the participation of that pathway in driving the conservation across the two model systems. B. The sankey diagram contains only the subset of cell line cancer (i.e., patient cancer pairs that have same tissue-specific lineage), and the cutoff for CS values is higher than 800 of the 1000 randomCSs obtained using the random selection of proteins.

### Pan-cancer stratification across model systems based on TransPRECISE scores

We deconvolved the global population-level networks to obtain sample-specific pathway-level functional summaries of the proteomic crosstalk within a pathway—in other words, for a given pathway, each sample has three different scores for activated, neutral, and suppressed statuses of the pathway. For the tumor stratification, we used the *network aberration score* defined as the sum of the activated and suppressed TransPRECISE scores for each sample.

For linking cell lines and patients, we computed the Pearson’s correlation for aberration score vectors (across twelve pathways) from each cell line-patient pair. Majority of the cell line-patient pairs for sarcoma-SARC (green), kidney-KIRC (light green), breast-BRCA (orange) and brain-LGG and GBM (light green and yellow) (edge colors in Figure 4 parenthesized) showed absolute correlations >0.9. Interestingly, the pancreatic and brain cancers were highly correlated across model systems: 99% of pancreas-HNSC pairs, 93% of GBM-pancreas pairs (and also 92% of the PAAD-head&neck pairs) had absolute correlations >0.9.

**Figure 4.**
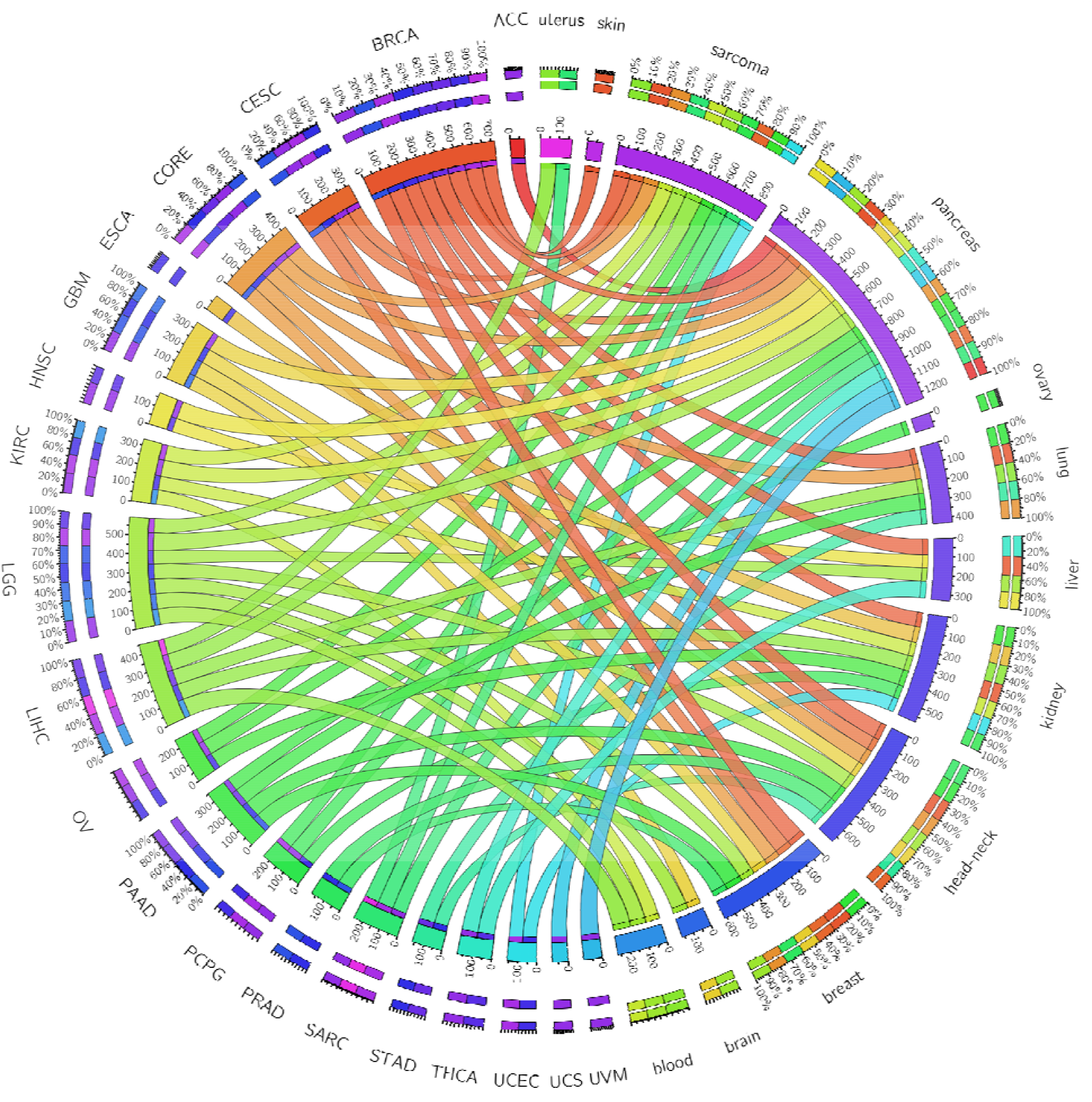
Circos plot summarizing high correlations of network aberration scores between patient and cell line cancers. An edge exists between a patient cancer type and a cell line cancer lineage if more than 75% of all possible patient-cell line pairs for that pair of cancers have a raw (Pearson) correlation of magnitude 0.9 or higher between their sets of the 12 pathway network aberration scores (sum of TransPRECISE sample-specific pathway activation and suppression scores). The edge strengths are determined by these percentages, as well. The edge colors indicate the patient cancers from which the edge originates, and the lengths of the innermost node pieces indicate the neighborhood size of the corresponding node. The two circular axes in the exterior indicate relative strengths of the edges originating from the same node, and the pieces here are colored by the opposite node to which that edge is connected, with the edges now arranged according to decreasing order of strength.

To find robust pan-cancer stratification across model systems, we applied hierarchical clustering using the complete linkage method^27^ on the correlations of the aberration scores. Among the 29 optimal clusters across patients and cell lines (Figure 5 and ST S8), most of the cell lines have mixed membership with patient tumors in 8 clusters (C2, C3, C4, C9, C13, C14, C19, and C23), while cluster C29 includes only cell lines (48 out of 640 in total, 7.5%). The cluster C4 showed a high level of fidelity in lineages between cell line and patient tumor types; it includes 81% of ovary cell lines and 11% of OV patients, 72% of head&neck cell lines and 38% of HNSC patients, and 20% of pancreas cell lines (another 70% of them being located in C2 with notable aberration of RAS/MAPK pathway), and 80% of PAAD patients exhibiting high aberration in apoptosis and DNA damage response pathways (ST S9). Within cluster C4, we observed significant correlations between the patient-cell line samples from ovary-PAAD, OV, BLCA, skin-PAAD, and head&neck-BLCA, HNSC (SF S11). More specifically, the HNSC samples were almost exclusively divided into the two clusters, C4 (n=78, 38%) and C15 (n=122, 60%), that include 38 (73%) head&neck cell lines and 5 (100%) oesophagus cell lines, respectively (Table S8). Patients with squamous cell carcinoma of the head and neck often have esophageal cancer.^28^ We found a significant difference in the survival outcome between HNSC patients in C4 and C15: the median survival was 456 days and 654 days for C4 and C15, respectively, with a p-value of 0.02 (Figure 5B). The patients in C15 that were represented by esophagus cell lines showed better survival than those in C4, which includes head&neck cell lines. This indicates that our TransPRECISE scores captured distinct prognostic information in HNSC patients. Moreover, the patterns of pathway activity and status were significantly different between the two clusters. The HNSC patients in C4 and C15 showed high aberration scores in apoptosis and PI3K/AKT pathways, respectively. While the aberration scores in the DNA damage pathway were high for both clusters, their PRECISE statuses were significantly distinct, with a Chi-squared test p-value <0.0001: 72% and 65% of the activated/suppressed patients in C4 and C15 showed suppression and activation, respectively.

**Figure 5.**
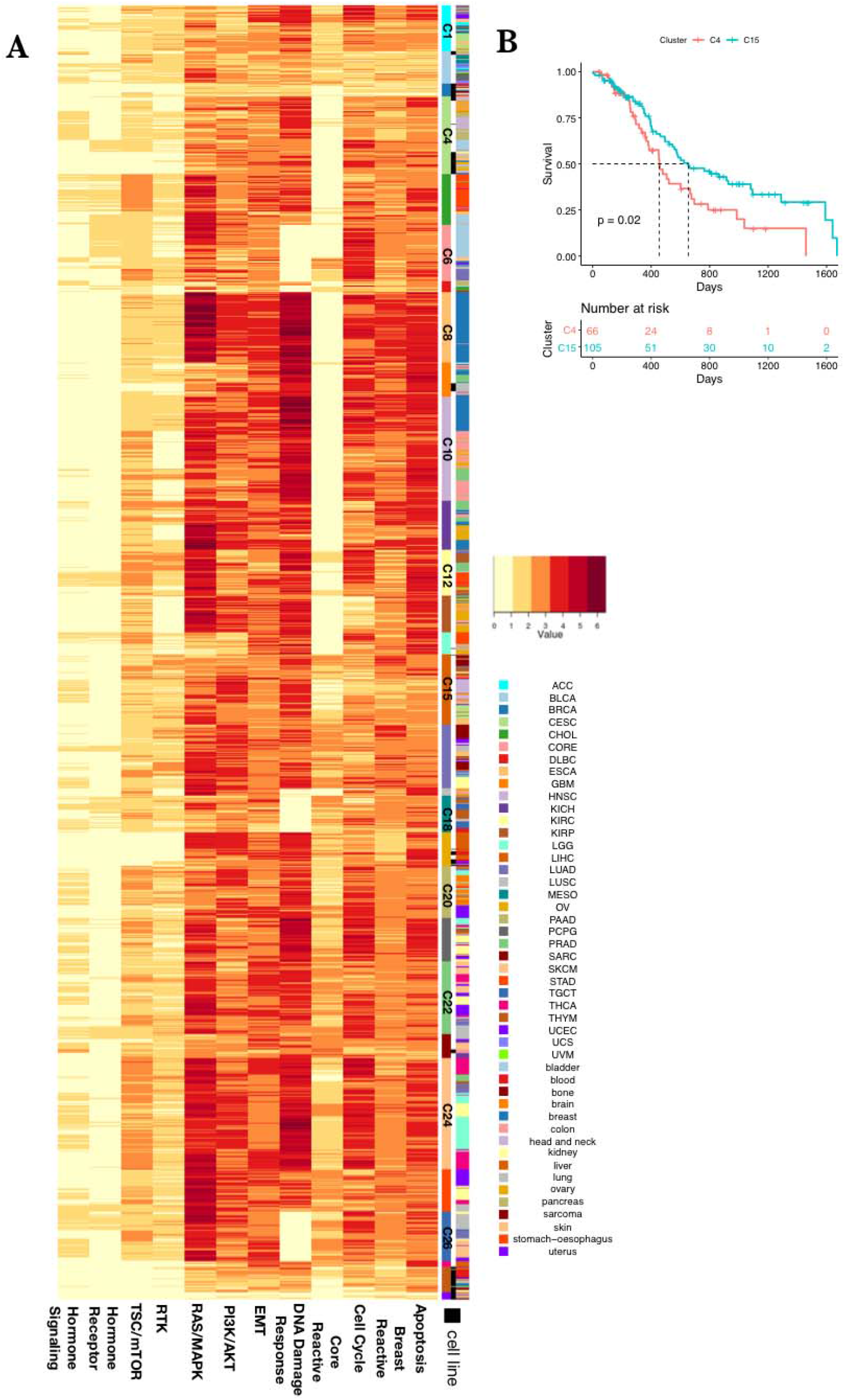
Avatar cell lines identification and selection of driving pathways using network aberration scores. A. Heatmap depicting network aberration scores (combined activation and suppression TransPRECISE pathway scores) after running unsupervised hierarchical clustering of the score matrix consisting of 8354 samples (7714 patients across 31 cancer lineages and 640 cell lines across 16 cancer types) and 12 proteomic signaling pathways. 29 clusters are identified by gap statistic, consisting of a mixture of patients and cell lines. Out of the three annotation bars, the topmost one indicates tumor types, the middle one indicates whether the sample is a patient or a cell line, and the bottom one indicates cluster participation according to which the samples are grouped. B. Kaplan-Meier plot depicting difference between survival times of HNSC patients that are clustered in clusters C4 and C15 using the hierarchical clustering method on TransPRECISE network aberration scores.

### Drug response prediction using TransPRECISE scores

#### Training drug response prediction models in cell lines

For the subset of cell lines where drug sensitivity data are available (ST S4), we used Bayesian additive regression trees (BART),^29^ a machine learning method, to build predictive models from the network aberration scores for the 12 pathways. For each cancer, we fit BART, with drug response (sensitive or resistant) as a binary outcome and TransPRECISE scores as predictors, for the drugs having profiles of ≥10 cell lines for that cancer type.

We found that TransPRECISE scores conferred high predictive power, translating to high median <u>test-set</u> areas under the receiver operating characteristic curves (AUCs) across the lineages; all lineages had median AUCs >0.8, with lung, breast, and colon being the top three, having median AUCs >0.9 (SF S12). From the radar plot summarizing the top pathway predictors across all drugs for each lineage (Figure 6A), we observed some notable evidence of predictive affinity for certain pathways to specific lineages: hormone receptor in breast, core reactive, RTK and TSC/mTOR in colon, RAS/MAPK in liver, DNA damage response and PI3K/AKT in lung, apoptosis, cell cycle and EMT in ovary, and DNA damage response and TSC/mTOR in pancreas cell lines. Further, we investigated pathway interaction in predicting drug sensitivity (Figure 6B and SF S13). The breast cancer related pathways, breast reactive, and hormone receptor pathways were highly synergistic in predicting the responses of five drugs including ML311 in breast cancer cell lines.^30^

**Figure 6.**
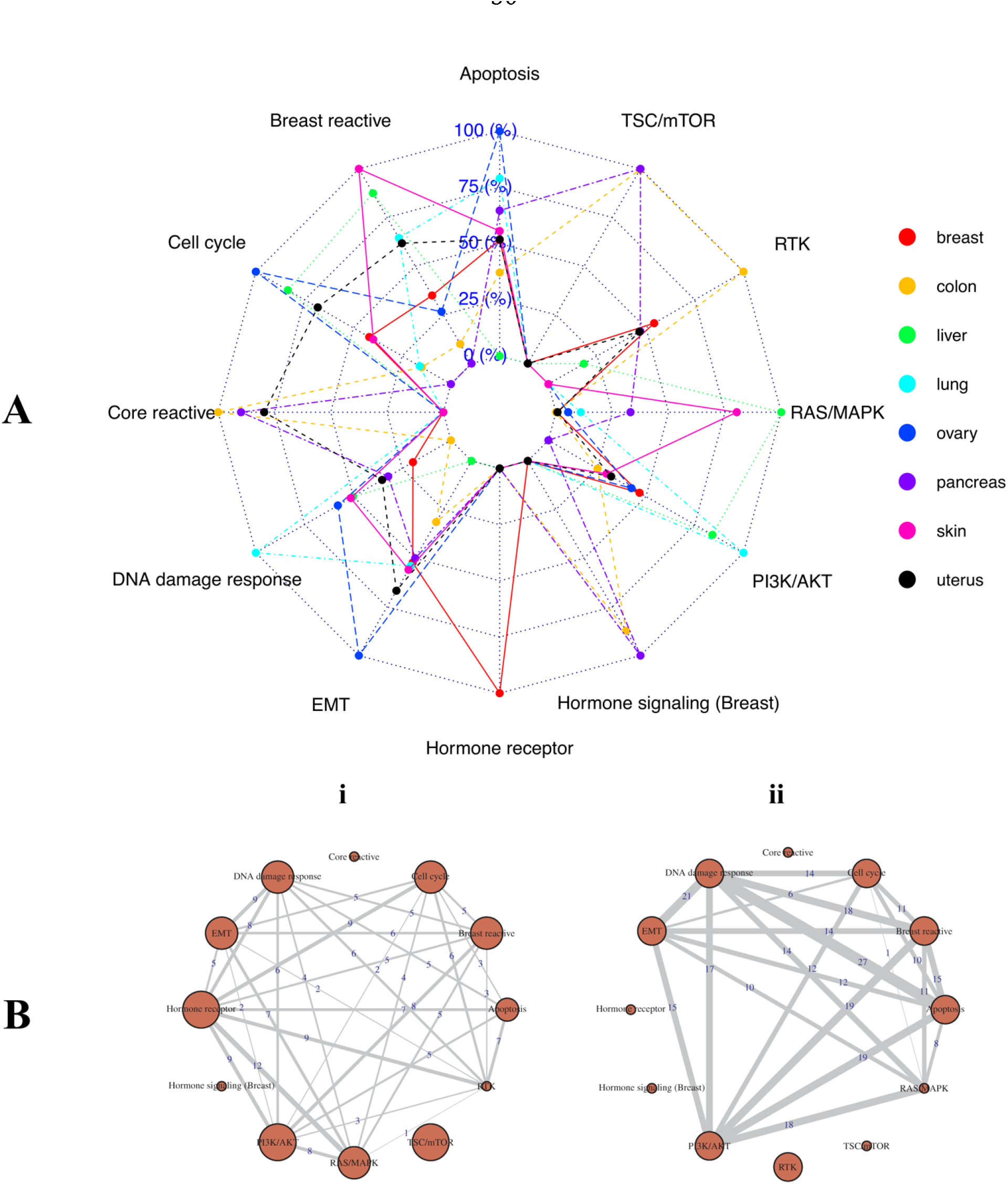
Performance of pathways in drug response prediction for cell lines across cancer lineages, based on test-set AUC values evaluated from fivefold cross validation. A. For a tissue type, we only look at the subset of drugs for which we have at least 10 response profiles from cell lines in that lineage and at least 0.85 test-set AUC using a five-fold cross-validation in the BART models. Then, for each pathway we compute the proportion of times it is the top predictor in models for such drugs. The radar plot shows these proportions in a log_e_ (1+.)- transformed scale. The significance and ranking of each of the twelve pathways in a model are quantified by posterior probabilities of inclusion in such a final predictive model for drugs. B. Networks showing the number of times (within models satisfying the criteria in panel A) a pair of pathways are the top two predictive pathways in a BART model. Panel i is for the breast cancer cell lines and panel ii is for the lung cancer cell lines.

#### Predicting drug sensitivity in patient tumors

For each of the cell line cancer lineage, for which the training models were fitted with the TransPRECISE pathway scores (as above), we predict drug sensitivity in patient tumors within matched tissue type (total 10 lineages). We found drugs that had 100% response rate especially in BRCA, CORE, LIHC, PAAD and SKCM; some of which are under clinical investigations in their respective cancers (ST S10 and SM Section S1.5). For example, all BRCA patients were predicted to be responsive to Ibrutinib that targets Bruton tyrosine kinase (BTK) with RAS/MAPK, PI3K/AKT, EMT as the top predictive pathways (ST S10).

## DISCUSSION

The investigation of patient tumors and cell-line interactome offers insights into the translational potential of preclinical model systems. This requires development of analytical models that capture the molecular heterogeneity of a cancer type in an unbiased manner and accurate calibration of aberrant biological pathways. We propose TransPRECISE, a multi-scale Bayesian network modeling framework, whose overarching goals are three-fold: identify differential and conserved intra-pathway activities between two different model systems (patient tumors and cell lines) across multiple cancers; globally assess cell lines as representative *in vitro* models for patients based on their inferred pathway circuitry; and build drug sensitivity prediction models for both cell lines and patients to aid pathway-based personalized medical decision-making. To the best of our knowledge, TransPRECISE is the first computational approach that provides a conflation of these goals.

As a proof-of-concept study, we illustrate the utility of TransPRECISE using RPPA-based proteomic expression profiles from patients and cell lines across several functional pathways, and a subset of the cell lines’ response to a number of drugs. At a pan-cancer level, we identified edges that were either (a) present in both the model systems with high prevalence, or, analogously, (b) present in only one of the model systems with prevalence. Type (a) edges offer valuable insights into the shared pathway circuitry across model systems, which has potential translational utility to study the role of tumor microenvironment. For example, category (a) edges included the CCNE2-FOXM1 edge in the cell-cycle pathway (10 cell line cancers, 17 patient cancers), which has been identified to have important implications in modulations of several cancers, such as breast,^31^ prostate cancer subtype 1,^32^ hepatocellular carcinoma,^33^ and osteosarcoma.^34^ At a cancer-specific level, we identified several {cell line, pathway, patient} triplets (based on network connectivity scores) that exhibited high degree of fidelity to their histological sites. A few specific tissue-pathway pairs identified by TransPRECISE analyses already have established translational value, including the RTK pathway in kidney cancers,^35,36^ and the hormone signaling pathway in ovarian cancers.^37,38^

As further validation, TransPRECISE implicated the EMT pathway in SKCM, which is expected since TCGA SKCM cohort contains many metastatic samples.^39^ EMT was also implicated in TCGA UCEC, another expected finding since UCEC includes epithelial-like endometrioid samples as well as mesenchymal-like serous samples in the cohort.^40^ Interestingly, TransPRECISE implicated the hormone receptor pathway in lung cancer, which is another known observation that is being studied for its translational potential.^41^ Those validation results increase our confidence in the findings from TransPRECISE.

The TransPRECISE-based sample-specific pathway scores can be used for identification of clinically relevant subtypes. Our robust pan-cancer stratification approach using the pathway scores provided subtypes within and across cancer that were represented by different cell line lineages (Figure 5) and contains distinct prognostic information. In particular, two different subsets of the head and neck cancer patients were included in two different clusters (C4 and C15), and had significantly different survival times, along with differing levels of activity in the DNA damage response pathway, potentially due to genomic aberrations in genes involved in that pathway.^42^

Using Bayesian machine learning models, we evaluated the prediction performance of TransPRECISE sample-specific pathway scores in predicting drug sensitivity in cell lines. A five-fold cross-validation study yielded test-set AUCs that confirmed the models’ high prediction accuracies across drugs (median AUC >0.8 for all cancers). Further investigations into the fitted-training models suggested notable pathway activities in predicting drug response in specific cancer tissues, such as hormone receptor-breast,^43^ and TSC/mTOR-pancreas.^44,45^ We extended our drug sensitivity analyses to the patient samples by using the training models from the matched cancer tissue to predict patients’ responses. Some of our findings, interestingly, are concordant with past and ongoing clinical trials. For example, ibrutinib, which had high predicted sensitivity for all the BRCA samples, has been investigated for its impact on HER2-amplified breast cancers.^46^ Similarly, lapatinib, in combination with trastuzumab, has recently been tested clinically for HER2-amplified metastatic colorectal cancer.^47^

One of the key strengths of the TransPRECISE algorithm is its generalizability, as it can be applied to any disease system that has matched genomic or molecular data on model and primary samples.

For example, the transition from RPPA to other advanced high-throughput platforms and development of databases, such as CPTAC,^48^ opens up the opportunity to include more proteins (thus, more pathways) in the network analyses: leading to a more global coverage of the proteomic crosstalk between model systems. Further, the PRECISE^26^ pipeline, which lies at the core of the TransPRECISE analyses, allows integration of upstream regulatory information and multi-omics layers such as mutations, copy number, methylation and mRNA expression. These modalities can be leveraged for better and holistic rewiring of pathway circuitry. Finally, our framework can be, in principle, applied to emerging model systems, such as patient-derived xenografts,^49,50^ and organoids,^51^ that allow better recapitulation of the human tumor microenvironment. In summary, TransPRECISE offers the potential to bridge the gap between human and pre-clinical models to delineate actionable cancer-pathway-drug interactions to assist personalized systems biomedicine approaches in the clinic.

## DATA AND MATERIAL AVAILABILITY

We have created an online, publicly available R shiny app (available at https://rupamb.shinyapps.io/transprecise) that is a comprehensive database and visualization repository of our findings. All codes used in generating our results are available, along with the documentation, on https://github.com/rupambh/TransPRECISE.

## ACKNOWLEDGEMENTS

This work was supported by the National Institutes of Health grants R21CA220299-01A1 (to M.J.H. and V.B.), U54-CA224065 (to M.J.H.) R01-CA160736, R01-CA194391, P30-CA46592 and National Science Foundation grant DMS 1922567 and funds from the UM Rogel Cancer Center and the School of Public Health to V.B; R01CA175486, U24CA209851, and U01CA217842, MD Anderson Faculty Scholar Award to H.L., and CCSG grant P30CA016672 (to H.L. and M.J.H.) and Cancer Prevention & Research Institute of Texas (CPRIT) grant RP180712 (to M.J.H.).

