## Supplementary Materials for "Personalized Network Modeling of the Pan-Cancer Patient and Cell Line Interactome"

### SECTION S1. SUPPLEMENTARY NOTES

#### Section S1.1 Pan-cancer multi-model systems data and preprocessing

Patient RPPA expression data We used reverse-phase protein array (RPPA) expression data for 7714 patients across 31 different cancer types available from The Cancer Proteome Atlas (TCPA) portal (Supplementary Table S1).[1,2]

Cell lines RPPA expression and drug sensitivity data We used RPPA expressions data for 640 cancer cell lines spanning across 16 different tissues (Supplementary Table S3) from the MD Anderson Cell Lines Project (MCLP).[3] Additionally, we used drug sensitivity data for a subset of size 254 out of the 640 cell lines with information on 481 drugs (Supplementary Table S4) from the Genomics of Drug Sensitivity in Cancer (GDSC)[4] database.

Imputing missing cell line expressions Unlike the patient tumor expression data, the cell lines expression data has some amount (approximately 6%) of missing values. We use the function "impute.knn" from the Bioconductor package "impute"[5] for imputation. For the k-nearest neighbor imputation implementation to run, each sample must have < 50% of missing data across all the variables, and each variable must have < 80% of missing data across all the samples. The original RPPA data set collected from MCLP has expression data on 651 cell lines from 194 genes that are common with our patient tumor RPPA dataset. Using the missing data upper bounds, we end up with 648 cell lines. We further removed eight more cell lines from some lineages (prostate (3), cervix (1), thyroid (1) and missing lineage (3)) that had too small sample sizes to be useful in fitting a stable Bayesian graphical regression model. We also removed one gene by the missing data criterion. Thus, we obtained our final set of 640 cell lines from 16 different cancer lineages with data on 193 proteins. We executed the imputation on this global profile consisting of all proteins at hand, instead of only the subset of proteins in the 12 functional pathways of interest

(Supplementary Table S4), since the imputation would be more informative this way and would not reflect any undue bias towards possible interactions within and between the pathways of interest. After completing imputation, we simply use the subset of the imputed data set with the proteins in the pathways of interest. The RPPA dataset for patients was used "as is," with the same subset of proteins selected, and was not put through any processing. For details on preprocessing of the drug sensitivity data on cell lines, please refer to Supplementary Materials Section S1.5.

**Section S1.2 TransPRECISE framework and model fitting steps**

Our Bayesian framework to estimate cancer-specific and sample-specific pathway networks and compute sample-specific pathway scores operates similarly to personalized cancer-specific integrated network estimation (PRECISE).[6] We use a Bayesian regression model, in which each protein is regressed on all the other proteins in the same pathway. The selected set of interacting protein-protein pairs then constitute the population-level cancer-specific pathway network. Given this population-level cancer-type-specific network, we deconvolve it to the sample-specific networks and scores, using the status of each protein (node) in the networks as neutral, suppressed, or activated. We then use the cancer-specific networks for pan-cancer and across model systems identification of conserved and differential pathway activities. We also use the sample-specific scores for identifying matching avatar cell lines for patient samples and predicting drug sensitivity.

*Step 1: Bayesian estimation of cancer-specific pathway networks* We aimed to estimate cancer-specific pathway networks using Bayesian regression methods on each of the proteins. We began with fixing one cancer type in one model system and one of the 12 pathways of interest. In this pathway of interest, for an interactive relationship among proteins, let $w_{ij} = w_{ji}$ be the weight connecting protein $i$ and protein $j$, with the number of proteins in the pathways denoted by $p$. We set $w_{ij} = w_{ji} = 0.5 \;\forall\; i \neq j$ and $w_{ii} = 0 \;\forall\; i$. Suppose $\mathrm{y_i}$ is the $n \times 1$ vector containing expression values of protein $i$ for $n$ samples from the fixed cancer lineage. For protein $i$, the $n \times 1$ expression vector $\mathrm{y_i}$ (centered with its mean) was modeled as

$$\mathrm{y_i} = \sum_{\mathrm{j \neq i}} \beta_{ij} y_j + \epsilon_\mathrm{i} = Z_\mathrm{i} \beta_\mathrm{i} + \epsilon_\mathrm{i},$$

where $\epsilon_\mathrm{i} \sim N_n(0_n, \sigma_i^2 I_n)$. $\{\beta_{ij} : j \neq i\}$ are the regression coefficients for the other proteins and $\beta_i$ is the vector of all of them. We employed Zellner's g-prior on $\beta_i$ as

$$\beta_i | g \sim N_{p-1}\left(0_{p-1}, \sigma_\mathrm{i}^2 \left(\frac{1}{g} Z_\mathrm{i}^T Z_\mathrm{i}\right)^{-1}\right).$$

The hyper-parameter $g$ reflects the prior on $\beta_i = 0$. A higher value of $g$ implies more probable deviation from $\beta_i = 0$, and we assigned $g = n$ by default, which yields the unit information prior in this case. We also set the prior $p(\sigma_i) \propto \sigma_i^{-1}$. Then we performed full enumeration using the Markov chain Monte Carlo algorithm. After performing all node-wise regressions following the above model, we select the median probability model to infer the posterior pathway network. Specifically, for each pair of proteins $i \; and \; j,$ the cancer-specific interaction between them is established if both $\beta_{ij} \; and \; \beta_{ji}$ correspond to posterior inclusion probabilities (defined as the sum of the posterior model probabilities for models where the covariate protein was included) > 0.5.

*Step 2: Construction of patient-specific networks via deconvolution* We further obtained a cancer-specific network with sample-specific labels on the nodes (proteins). Specifically, the activation statuses of the nodes are evaluated by estimating the posterior predictive density for each protein for each sample. To determine the activation status of a protein $i$ for a sample $j$ ($y_{ij}$), we computed the posterior probabilities of the protein expression to lie in the $\delta$-interval around zero ($p_{ij}^0$), to be greater than $\delta$ ($p_{ij}^+$), or less than $-\delta$ ($p_{ij}^-$). Then, we decided whether a protein is neutral, activated, or suppressed, depending on the maximum of these three posterior probabilities. Thus, samples from the same cancer lineage may have different node labels as suppressed, neutral, or activated, while the structure of the networks stay same across all such samples. Using $\delta = 0.5$, we calculated TransPRECISE networks across all samples for each of the 31 patient cancers and 16 cell line cancers, and each of the 12 pathways.

<u>*Step 3: Calibrating patient-specific pathway scores and status*</u> To compute an aggregated pathway activity score for each sample, we derive summary measures from the TransPRECISE networks obtained from previous step, indicating the entire pathway as neutral, activated, or suppressed. Under the TransPRECISE networks, the number of nodes that are connected to protein $i$, $(j : i \leftrightarrow j\,)$ is denoted by $C_i$. For a given pathway with $p$ genes, the pathway activity scores for a sample $j$ are given by the following set of equations for activated, suppressed, and neutral TransPRECISE scores, respectively.

$$\kappa_j^+ := \frac{1}{p}\sum_{i=1}^{p} p_{ij}^+(C_i + 1),\ \kappa_j^- := \frac{1}{p}\sum_{i=1}^{p} p_{ij}^-(C_i + 1), \text{ and } \kappa_j^0 := \frac{1}{p}\sum_{i=1}^{p} p_{ij}^0(C_i + 1).$$

Note that these sample-specific pathway scores are weighted averages of the posterior probabilities for suppressed ($p_{ij}^-$), neutral ($p_{ij}^0$), and activated ($p_{ij}^+$) statuses of the proteins by the number of connected proteins. Therefore, hub proteins in the pathway that exercise more control over the network through higher number of interacting proteins get higher weights towards determining the cumulative network score. For a given pathway and each sample, the *TransPRECISE pathway status*—which indicates whether the pathway is suppressed, neutral, or activated for the sample—is then decided by maximum of the three TransPRECISE pathway scores (i.e., max $\{\kappa_j^+, \kappa_j^-, \kappa_j^0\}$) for each sample $j$.

**Section S1.3 Cancer-specific protein networks and the rewiring across pathways**

<u>*Comparing network activity levels across model systems*</u> For a given cancer lineage with respect to a pathway, the connectivity score (CS) is the ratio of the observed number of edges in the cancer-specific network to the total number of possible edges in the pathway ($\frac{p(p-1)}{2}$, if the pathway has

$p$ proteins). The suggestion of significant interaction exhibited within a pathway for a cancer type is then quantified by finding a randomCS proportion based on the CS, a low value of which indicates higher nonrandom interactions in the pathway for that cancer type. Briefly, for each cancer type and pathway, we randomly select the same number of proteins as in that given pathway from the pool of all proteins across the 12 pathways. After constructing TransPRECISE networks from a total of 1000 such random permutations of the proteins, a 1000 new values of the CS (called randomCS, which would be coming from the underlying null distribution if there were no significant interaction within the pathway, in addition to the possible interactions already present in the global pool of proteins) for that cancer type and pathway are obtained. The randomCS proportion corresponding to this cancer type and pathway pair is then defined by the following equation:

$$p_{CS} := \frac{\sum_{i=1}^{1000} I[CS_i > CS]}{1000}.$$

Here $I(.)$ is an indicator function, $CS_i$ is the randomCS value obtained from the i$^{th}$ permutation and $CS$ is the actual CS value obtained from the data. Now, suppose we have a patient cancer type $P$ and a cell line cancer type $C$, say. For a pathway $G$ fixed, we say that the triplet $P - G - C$ is connected in terms of similar network activity, if we have both $p_{CS_{P,G}}$ and $p_{CS_{C,G}} < \epsilon$ for some small chosen value of $\epsilon$. Figure 3A then depicts all such triplets for $\epsilon = 0.1$, and Figure 3B depicts all such triplets where $P\ and\ C$ come from the same tissue for $\epsilon = 0.2$. A higher value of $\epsilon$ would result in a higher number of connections and decreasing the cutoff would lead to more refined sets of edges that have strong suggestion towards conserved network activity. The cutoffs used were chosen empirically by checking at what magnitude of the cutoff the number of connections in the corresponding networks have a large decrease.

**Section S1.4 Correlating patient and cell line tumors based on PRECISE scores**

*Construction of pathway network aberration scores* For a given pathway, the network aberration score of a sample $j$ is defined as $\kappa_j^+ + \kappa_j^-$, where $\kappa_j^+$ and $\kappa_j^-$ are the respective activated and suppressed TransPRECISE scores, as defined in Section S1.2.

*Hierarchical clustering of patients and cell lines* Using the resulting TransPRECISE network aberration score matrix as the input data, we obtain a robust pan-cancer and pan-model systems stratification. The data matrix has 8354 rows, the first 7714 of them corresponding to one patient each and the next 640 corresponding to one cell line each, and 12 columns, each corresponding to a pathway. Based on the Euclidean distance of the score matrix, we applied hierarchical clustering using Ward's method.[7] To determine the number of clusters, we used the gap statistic.[8]

**Section S1.5 Predicting drug response using PRECISE scores**

*Training models for cell lines' drug response* Out of the 254 cell lines that have drug sensitivity information, eight cancer lineages can be obtained with at least 10 samples (Supplementary Table S4). For each lineage and for each drug which has at least 10 non-missing responses for that lineage, we fit a Bayesian additive regression tree (BART)[9] model using the package "bartMachine",[10] with the predictors being the 12 pathway network aberration scores and the response variable being the binary (sensitive/resistant) drug response. We compute the area under the receiver operating characteristics curve (AUC) for each model with a fivefold cross-validation, and only keep those models for further inferences that had test-set AUC >0.85. We obtain the ranking of predictors in each model using the variable inclusion proportions, defined as the number of times a covariate is selected in a model within the 1000 iterations after a 250-iteration burn-in.

*Predicting patient drug responses* For each of the eight cell line lineages with at least 10 samples in the drug sensitivity data, we use the corresponding models for predicting the drug sensitivity in patient tumors with the same tissue type. The predictions are in a continuous scale of 0-1, and we mark a sample as “sensitive” if the predicted response is >0.5. We define the ‘response rate’ of a patient cancer with respect to a drug as the proportion of samples labeled sensitive within each for that cancer-drug combination and rank the drugs within each cancer type in decreasing order of these response rates.

### SECTION S2. SUPPLEMENTARY FIGURES AND TABLES

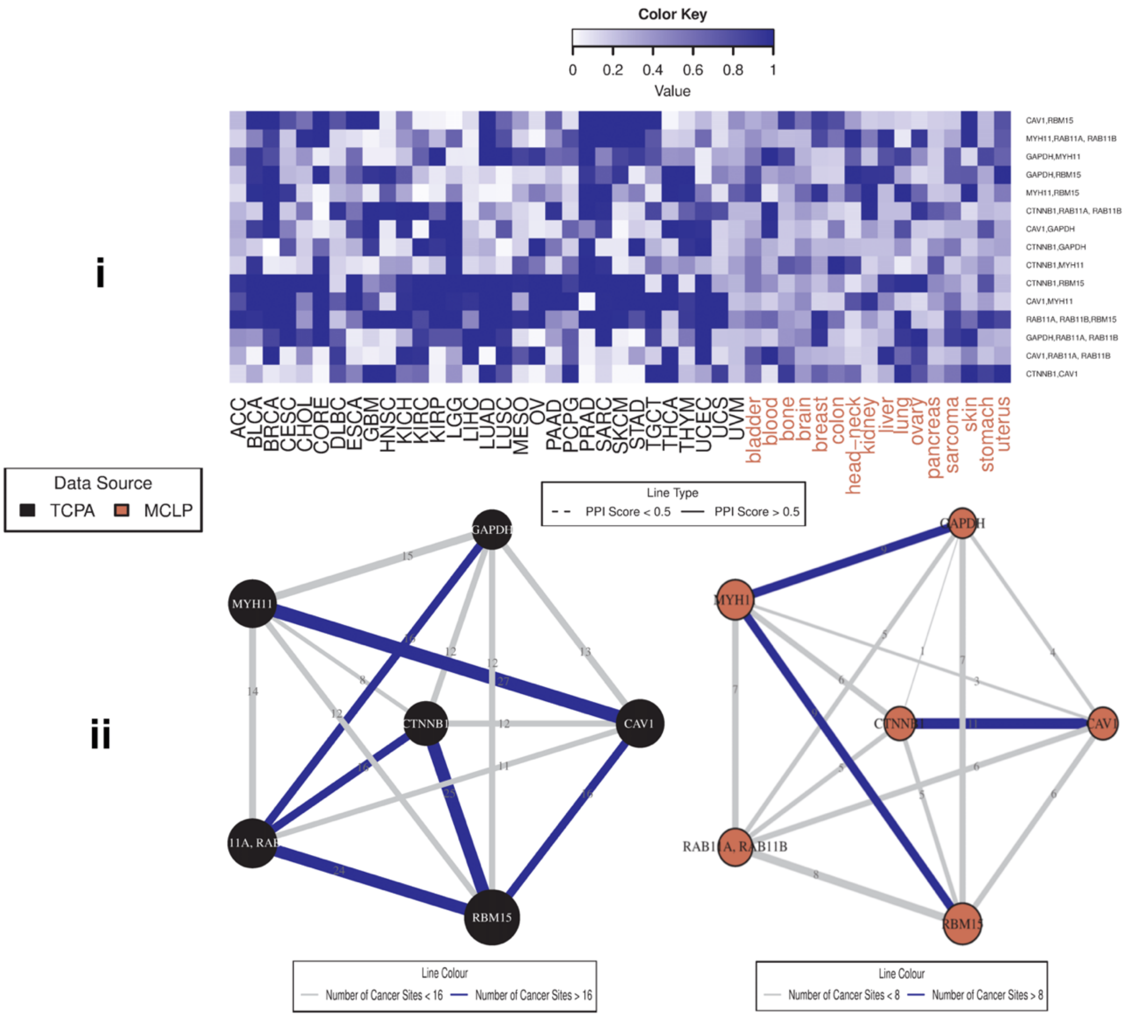

**Supplementary Figure S1. Pan-cancer summary of protein networks for breast reactive pathway.** i. Heatmap depicting strengths of all possible protein-protein edges within the pathway, across all 47 patient and cell line tumor lineages, quantified by the posterior inclusion probabilities of the edges based on the fitted Bayesian graphical regression model. ii. Left panel exhibits a network with its edges weighted and labeled by the edge consistencies (ECs), which are quantified by the number of patient tumor types holding that particular edge, also presenting the *a priori* known strength of the edge using the protein-protein interaction score from the STRING database. The right panel is the corresponding network across cell line cancers.

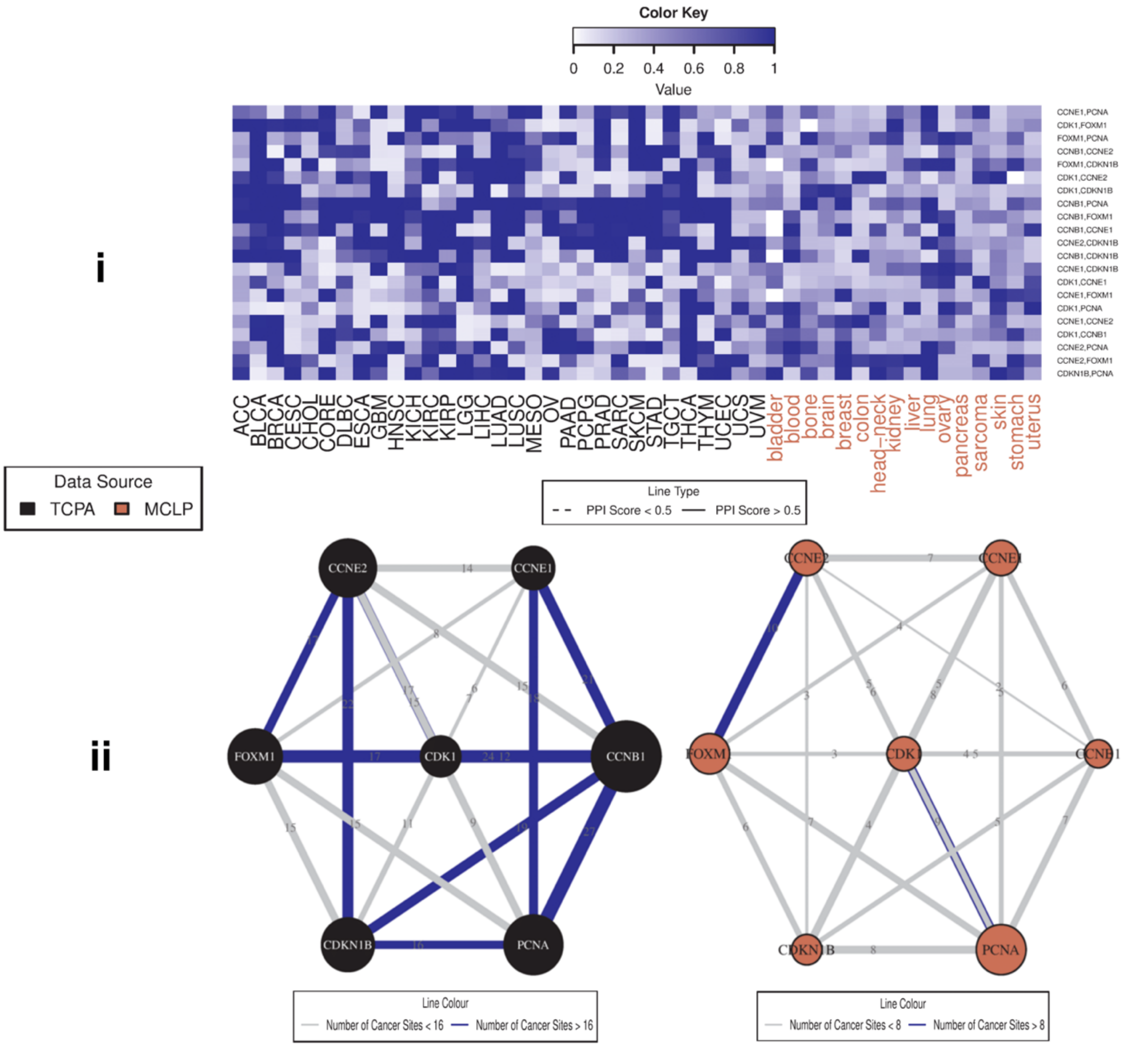

**Supplementary Figure S2. Pan-cancer summary of protein networks for cell cycle pathway.** i. Heatmap depicting strengths of all possible protein-protein edges within the pathway, across all 47 patient and cell line tumor lineages, quantified by the posterior inclusion probabilities of the edges based on the fitted Bayesian graphical regression model. ii. Left panel exhibits a network with its edges weighted and labeled by the edge consistencies (ECs), which are quantified by the number of patient tumor types holding that particular edge, also presenting the *a priori* known strength of the edge using the protein-protein interaction score from the STRING database. The right panel is the corresponding network across cell line cancers.

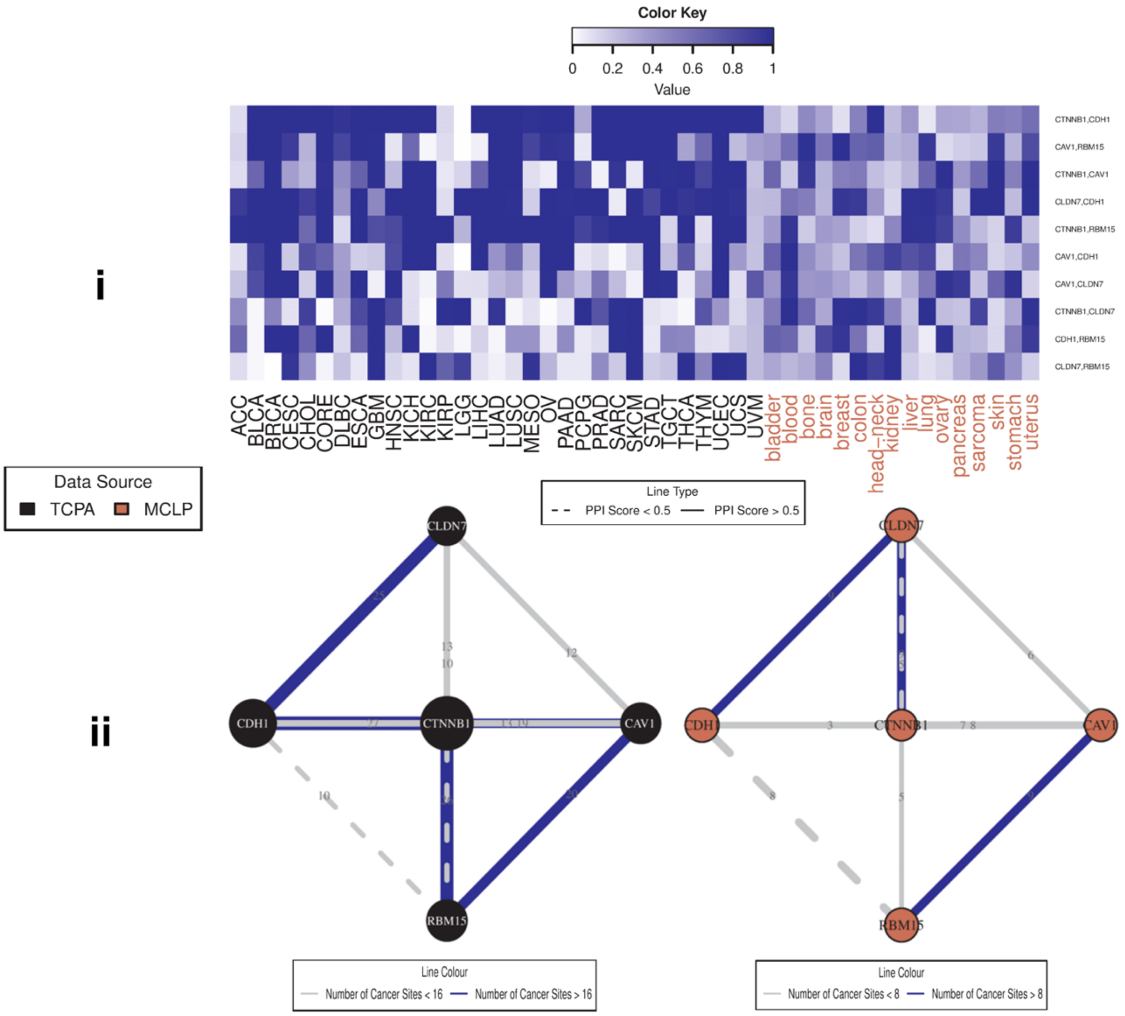

**Supplementary Figure S3. Pan-cancer summary of protein networks for core reactive pathway.** i. Heatmap depicting strengths of all possible protein-protein edges within the pathway, across all 47 patient and cell line tumor lineages, quantified by the posterior inclusion probabilities of the edges based on the fitted Bayesian graphical regression model. ii. Left panel exhibits a network with its edges weighted and labeled by the edge consistencies (ECs), which are quantified by the number of patient tumor types holding that particular edge, also presenting the *a priori* known strength of the edge using the protein-protein interaction score from the STRING database. The right panel is the corresponding network across cell line cancers.

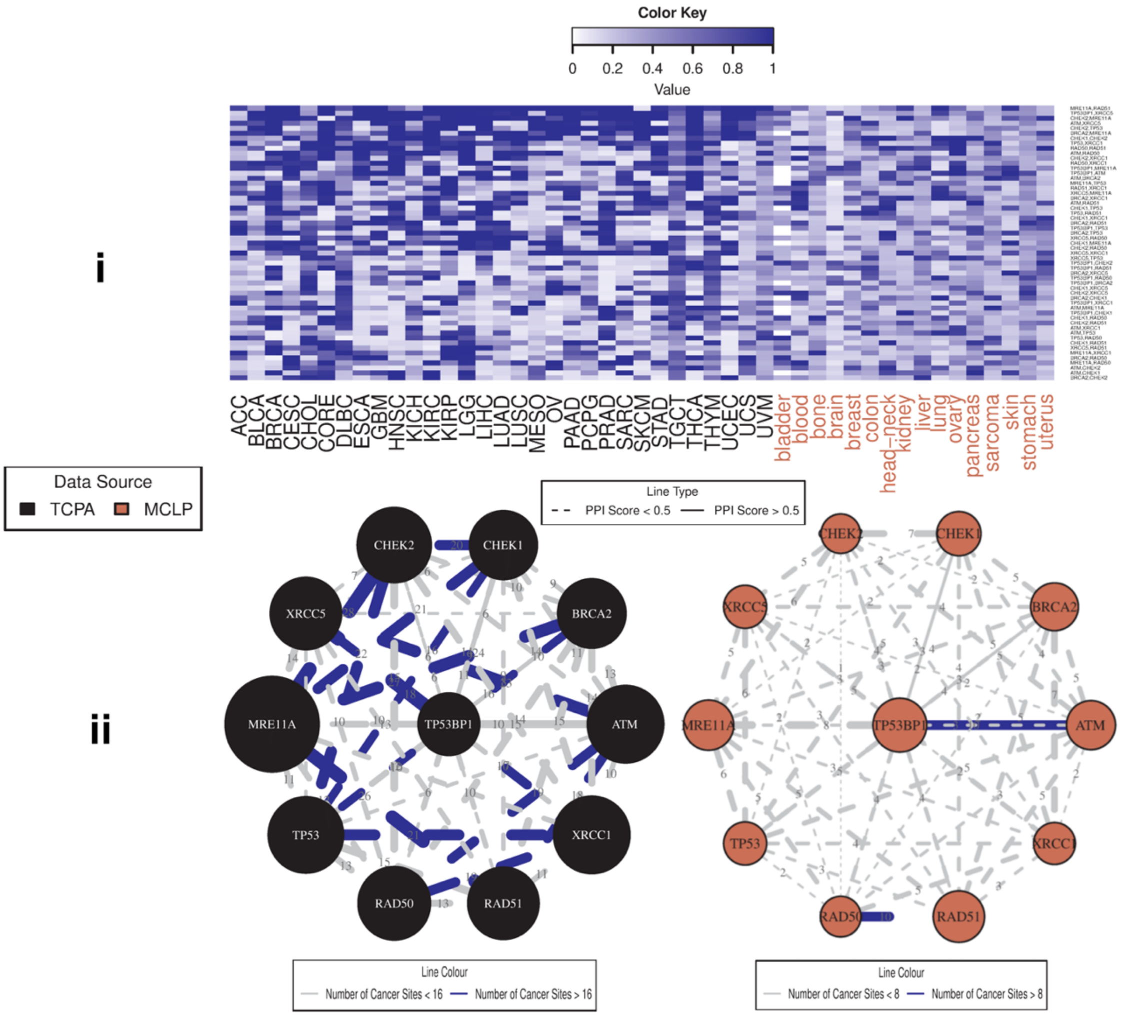

**Supplementary Figure S4. Pan-cancer summary of protein networks for DNA damage response pathway.** i. Heatmap depicting strengths of all possible protein-protein edges within the pathway, across all 47 patient and cell line tumor lineages, quantified by the posterior inclusion probabilities of the edges based on the fitted Bayesian graphical regression model. ii. Left panel exhibits a network with its edges weighted and labeled by the edge consistencies (ECs), which are quantified by the number of patient tumor types holding that particular edge, also presenting the *a priori* known strength of the edge using the protein-protein interaction score from the STRING database. The right panel is the corresponding network across cell line cancers.

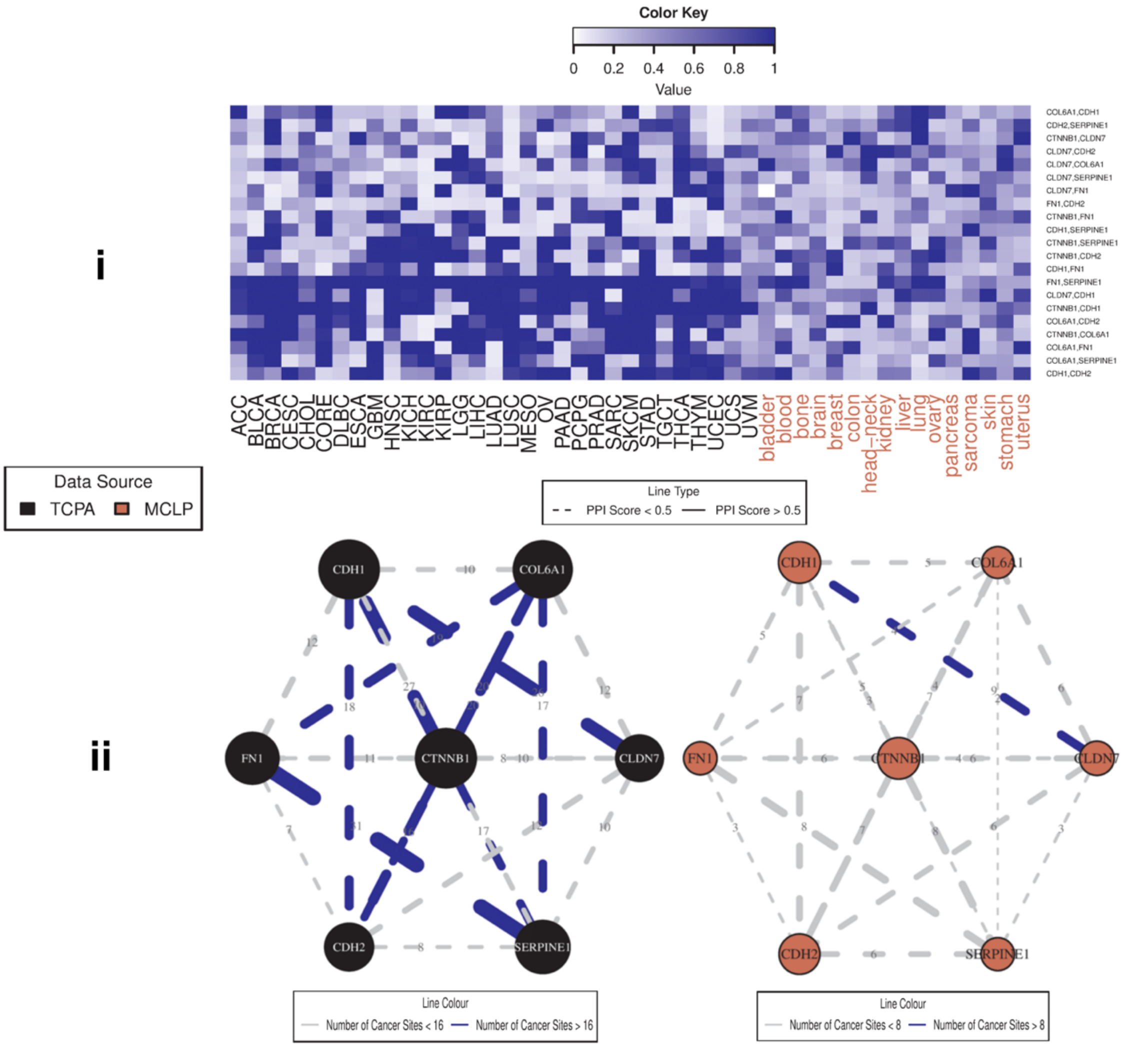

**Supplementary Figure S5. Pan-cancer summary of protein networks for EMT pathway.** i. Heatmap depicting strengths of all possible protein-protein edges within the pathway, across all 47 patient and cell line tumor lineages, quantified by the posterior inclusion probabilities of the edges based on the fitted Bayesian graphical regression model. ii. Left panel exhibits a network with its edges weighted and labeled by the edge consistencies (ECs), which are quantified by the number of patient tumor types holding that particular edge, also presenting the *a priori* known strength of the edge using the protein-protein interaction score from the STRING database. The right panel is the corresponding network across cell line cancers.

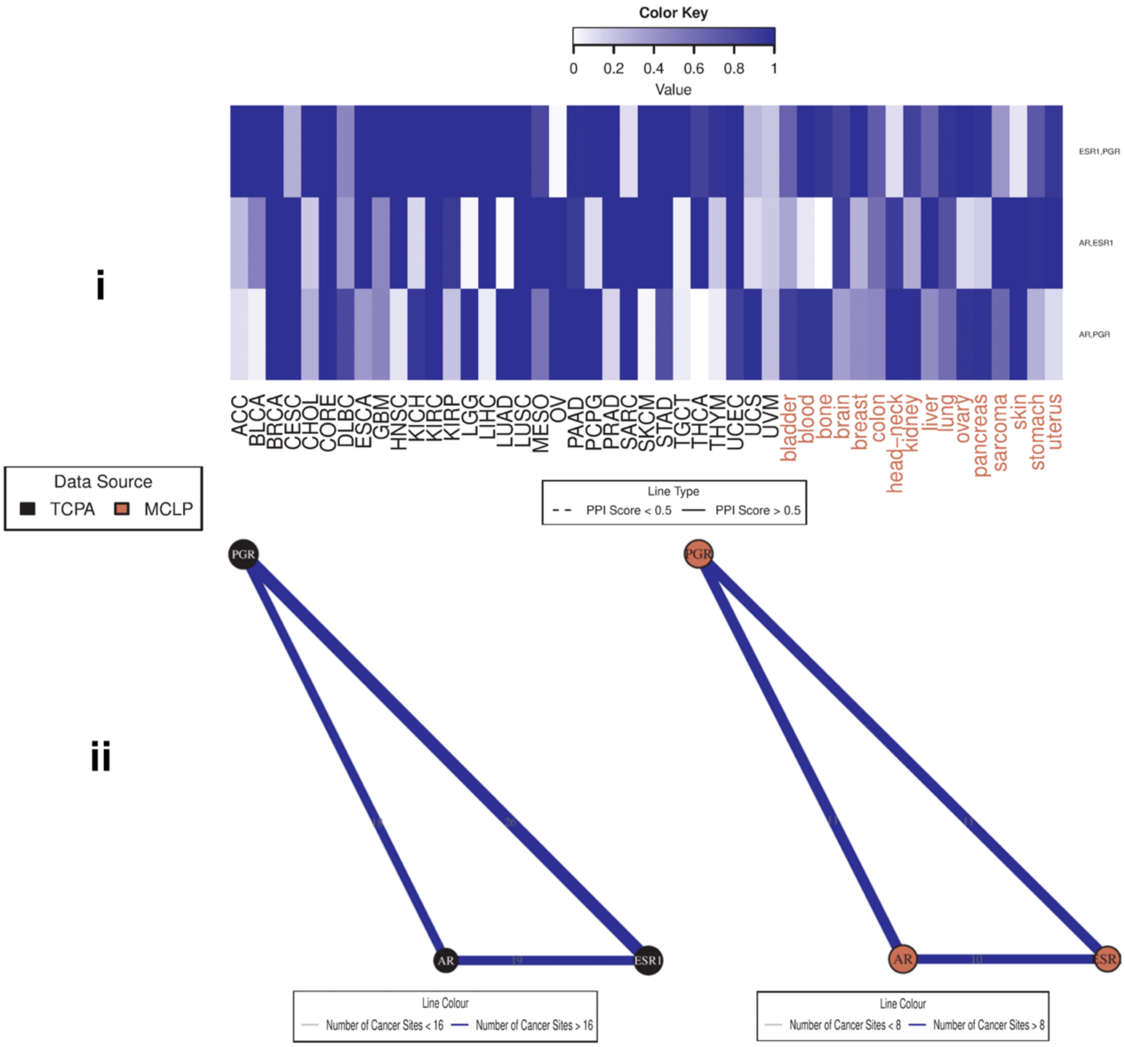

**Supplementary Figure S6. Pan-cancer summary of protein networks for hormone receptor pathway.** i. Heatmap depicting strengths of all possible protein-protein edges within the pathway, across all 47 patient and cell line tumor lineages, quantified by the posterior inclusion probabilities of the edges based on the fitted Bayesian graphical regression model. ii. Left panel exhibits a network with its edges weighted and labeled by the edge consistencies (ECs), which are quantified by the number of patient tumor types holding that particular edge, also presenting the *a priori* known strength of the edge using the protein-protein interaction score from the STRING database. The right panel is the corresponding network across cell line cancers.

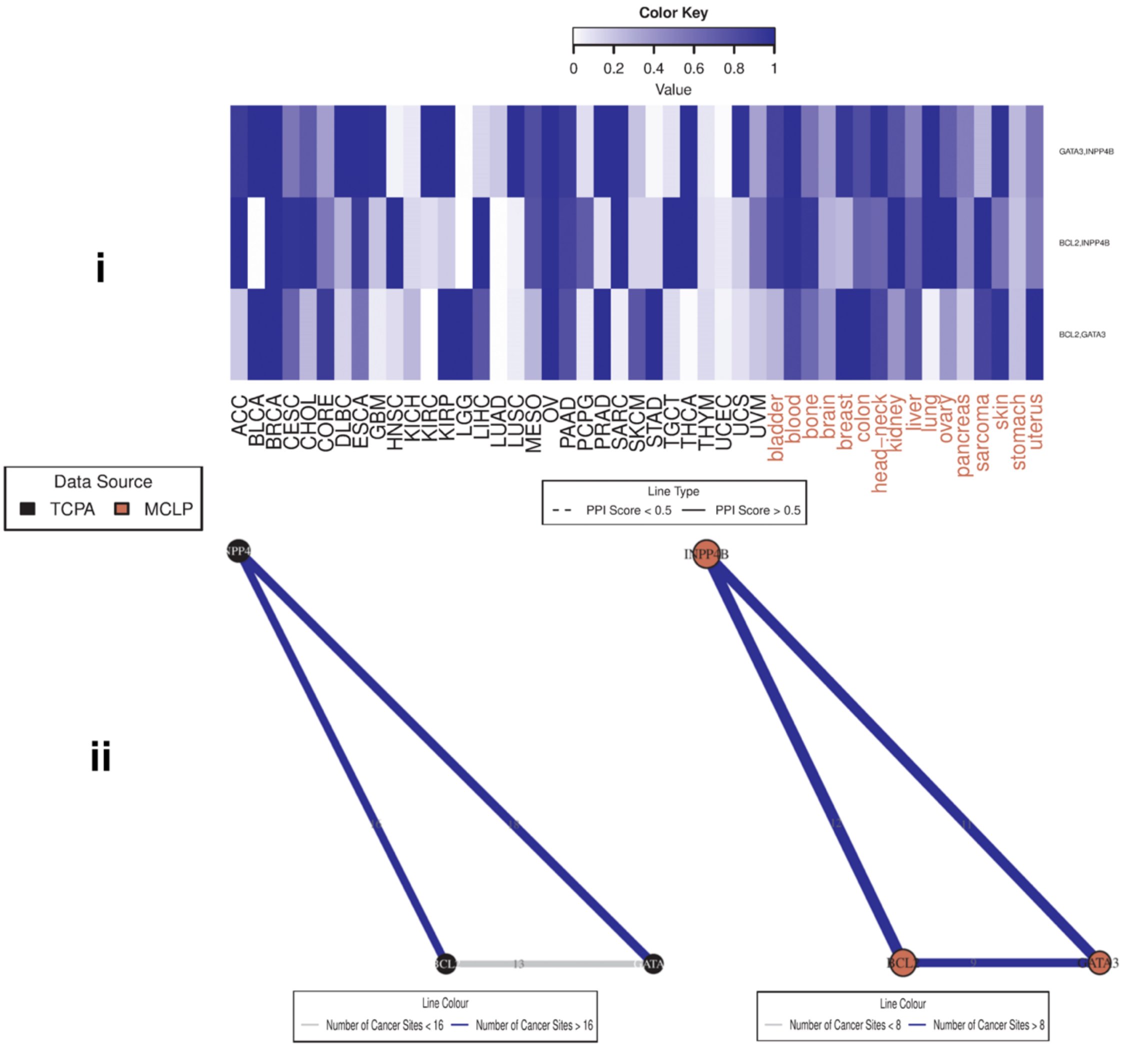

**Supplementary Figure S7. Pan-cancer summary of protein networks for hormone signaling (breast) pathway.** i. Heatmap depicting strengths of all possible protein-protein edges within the pathway, across all 47 patient and cell line tumor lineages, quantified by the posterior inclusion probabilities of the edges based on the fitted Bayesian graphical regression model. ii. Left panel exhibits a network with its edges weighted and labeled by the edge consistencies (ECs), which are quantified by the number of patient tumor types holding that particular edge, also presenting the *a priori* known strength of the edge using the protein-protein interaction score from the STRING database. The right panel is the corresponding network across cell line cancers.

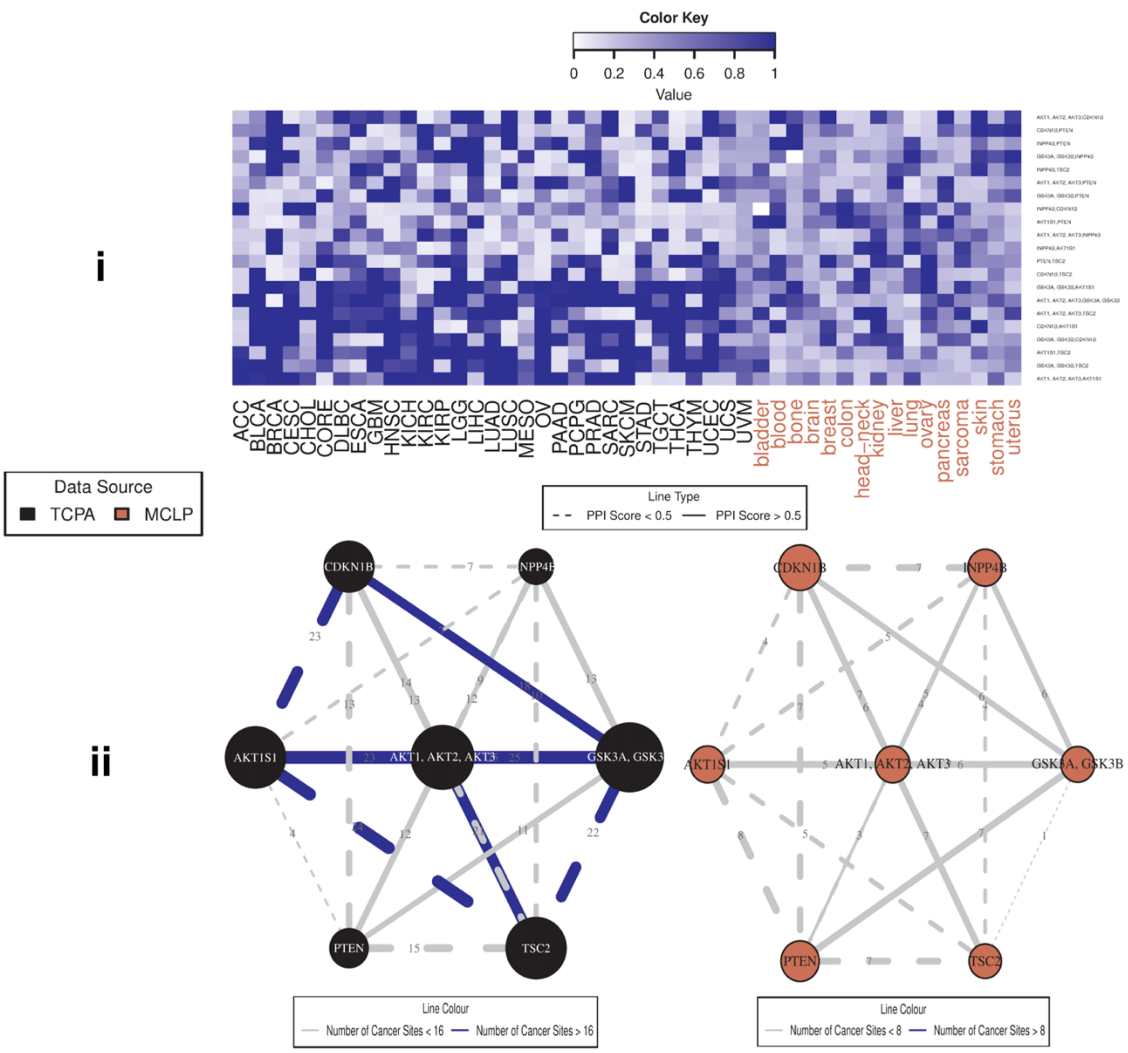

**Supplementary Figure S8. Pan-cancer summary of protein networks for PI3K/AKT pathway.** i. Heatmap depicting strengths of all possible protein-protein edges within the pathway, across all 47 patient and cell line tumor lineages, quantified by the posterior inclusion probabilities of the edges based on the fitted Bayesian graphical regression model. ii. Left panel exhibits a network with its edges weighted and labeled by the edge consistencies (ECs), which are quantified by the number of patient tumor types holding that particular edge, also presenting the *a priori* known strength of the edge using the protein-protein interaction score from the STRING database. The right panel is the corresponding network across cell line cancers.

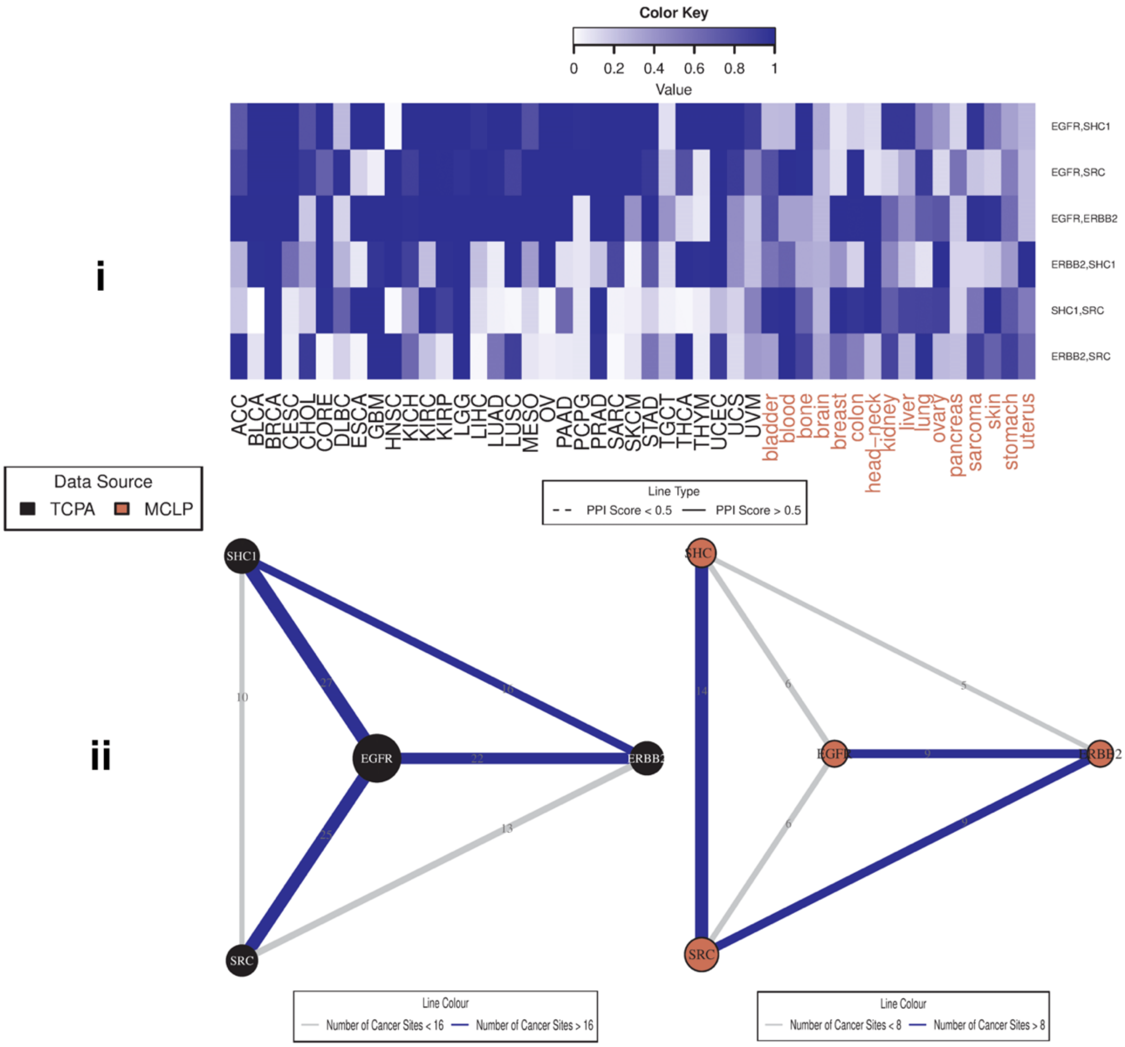

**Supplementary Figure S9. Pan-cancer summary of protein networks for RTK pathway.** i. Heatmap depicting strengths of all possible protein-protein edges within the pathway, across all 47 patient and cell line tumor lineages, quantified by the posterior inclusion probabilities of the edges based on the fitted Bayesian graphical regression model. ii. Left panel exhibits a network with its edges weighted and labeled by the edge consistencies (ECs), which are quantified by the number of patient tumor types holding that particular edge, also presenting the *a priori* known strength of the edge using the protein-protein interaction score from the STRING database. The right panel is the corresponding network across cell line cancers.

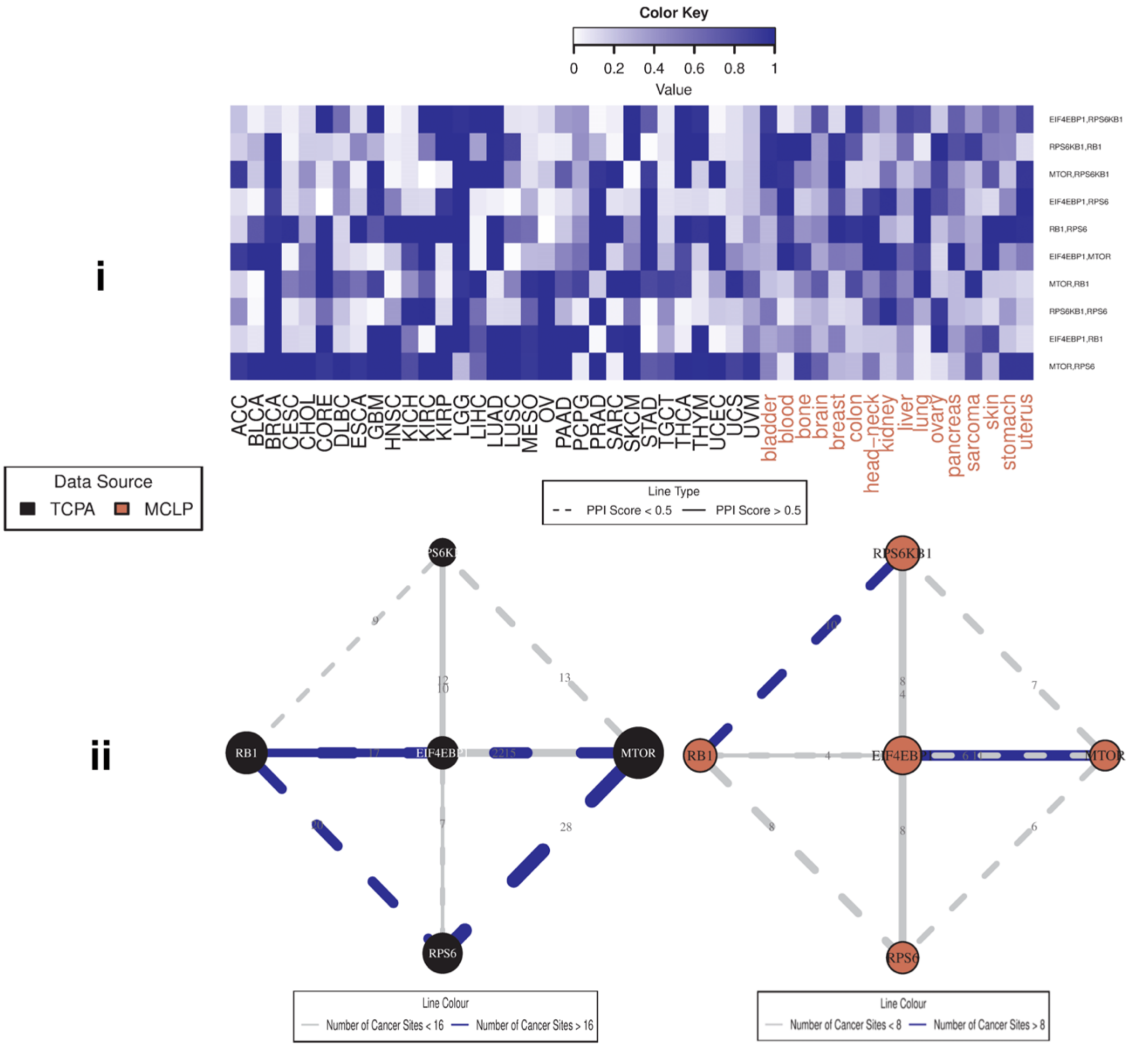

**Supplementary Figure S10. Pan-cancer summary of protein networks for TSC/mTOR pathway.** i. Heatmap depicting strengths of all possible protein-protein edges within the pathway, across all 47 patient and cell line tumor lineages, quantified by the posterior inclusion probabilities of the edges based on the fitted Bayesian graphical regression model. ii. Left panel exhibits a network with its edges weighted and labeled by the edge consistencies (ECs), which are quantified by the number of patient tumor types holding that particular edge, also presenting the *a priori* known strength of the edge using the protein-protein interaction score from the STRING database. The right panel is the corresponding network across cell line cancers.

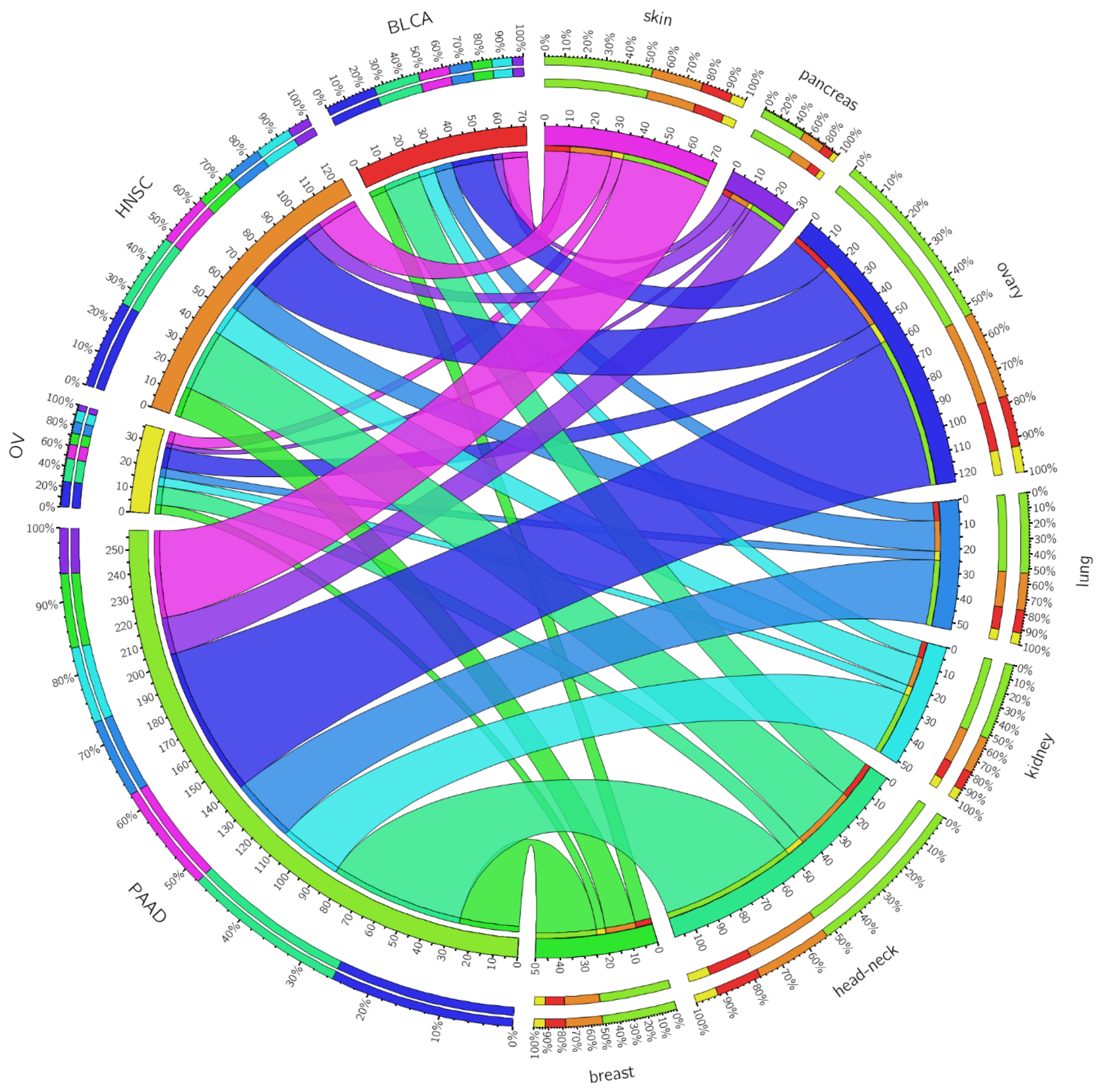

**Supplementary Figure S11. Circos plot summarizing joint membership of patient and cell line samples in cluster C4.** An edge exists between a patient cancer type and a cell line cancer lineage if more than 10% of total number of samples for both of the lineages are located in cluster C4. The edge strengths are determined by the product of the two percentages for the two nodes (lineages) scaled by 100. The edge colors pertain to the cell line cancers that the edge originates from, and the lengths of the innermost node pieces indicate the neighborhood size of the corresponding node. The two circular axes in the exterior indicate relative strengths of the edges originating from the same node, and the pieces here are colored by the opposite node to which that edge is connected, with the edges now arranged according to decreasing order of strength.

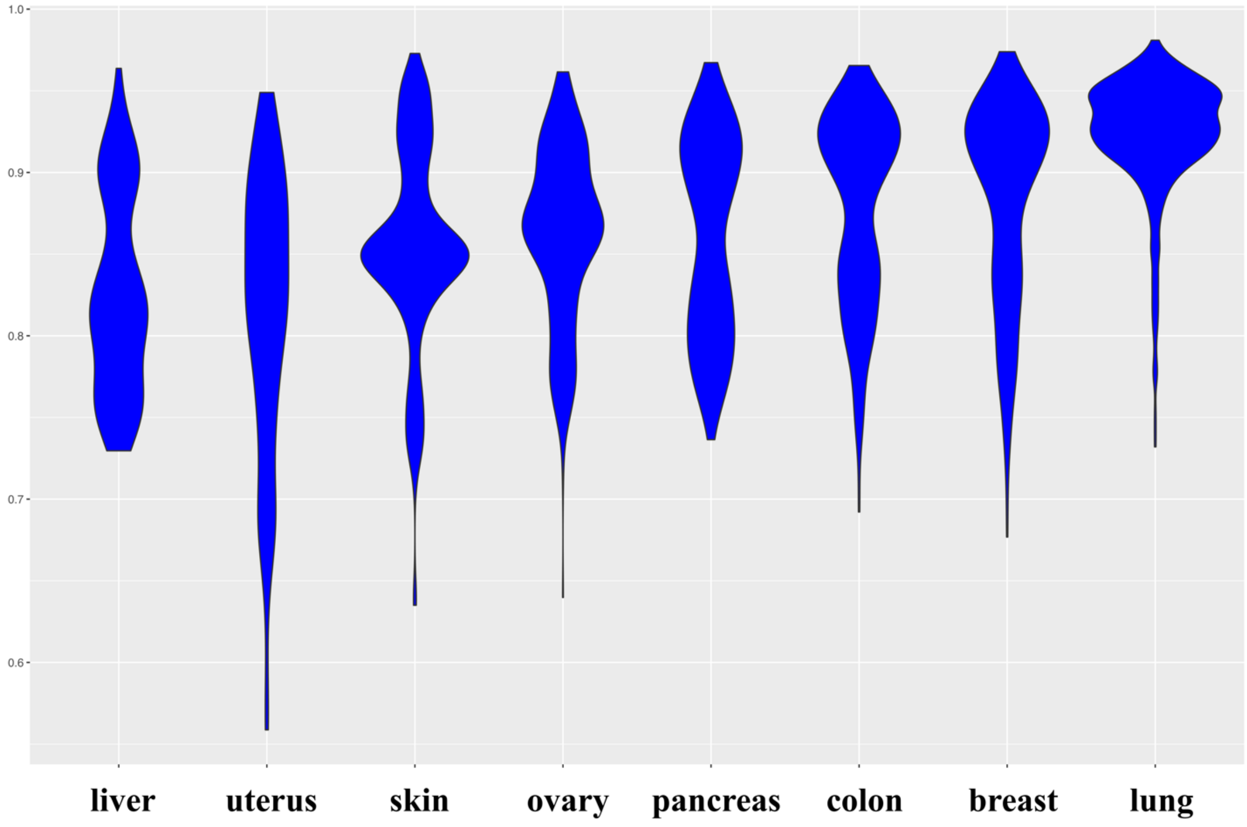

**Supplementary Figure S12. Performance of drug sensitivity prediction models across cell line lineages.** Each column exhibits a violin plot for AUCs from all the fitted BART models for the corresponding cancer type for each drug with at least 10 responses available for the samples in that lineage.

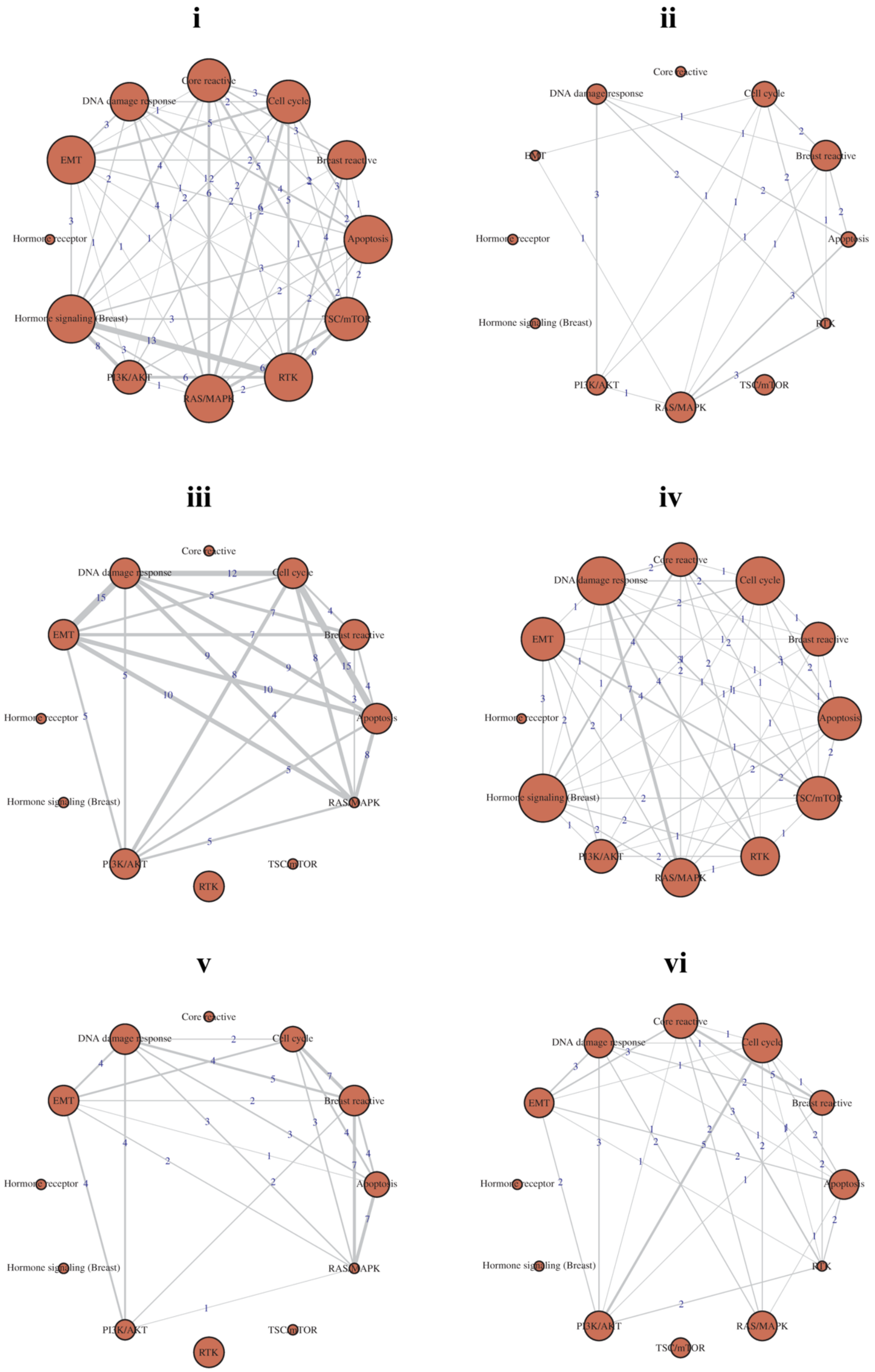
i
Core reactive
DNA damage response
Cell cycle
EMT
Breast reactive
Hormone receptor
Apoptosis
Hormone signaling (Breast)
TSC/mTOR
PI3K/AKT
RAS/MAPK
RTK
ii
iii
iv
v
vi

**Supplementary Figure S13. Shared presence of pathways in cell line drug sensitivity prediction models.** For each lineage, we only look at drugs with at least 10 response profiles available within cell lines from that lineage and the corresponding BART model having a >0.85 test-set AUC based on five-fold cross-validations. Edge weights indicate the number of times the two nodes (pathways) are the top two predictors within all such BART models for a lineage. Each panel corresponds to a different lineage, as follows: i. colon, ii. liver, iii. ovary, iv. pancreas, v. skin, vi. uterus.

**Supplementary Table S1.** Summary of patient tumor sample sizes according to lineages.

| Cancer Lineage | Study Abbreviation | Number of Patients |
|---|---|---|
| Adrenocortical carcinoma | ACC | 46 |
| Bladder Urothelial Carcinoma | BLCA | 343 |
| Breast invasive carcinoma | BRCA | 878 |
| Cervical squamous cell carcinoma and endocervical adenocarcinoma | CESC | 171 |
| Cholangiocarcinoma | CHOL | 30 |
| Colon/Rectum adenocarcinoma | CORE | 491 |
| Lymphoid Neoplasm Diffuse Large B-cell Lymphoma | DLBC | 33 |
| Esophageal carcinoma | ESCA | 126 |
| Glioblastoma multiforme | GBM | 232 |
| Head and Neck squamous cell carcinoma | HNSC | 203 |
| Kidney Chromophobe | KICH | 63 |
| Kidney renal clear cell carcinoma | KIRC | 469 |
| Kidney renal papillary cell carcinoma | KIRP | 217 |
| Brain Lower Grade Glioma | LGG | 432 |
| Liver hepatocellular carcinoma | LIHC | 184 |
| Lung adenocarcinoma | LUAD | 362 |
| Lung squamous cell carcinoma | LUSC | 325 |
| Mesothelioma | MESO | 61 |
| Ovarian serous cystadenocarcinoma | OV | 431 |
| Pancreatic adenocarcinoma | PAAD | 122 |
| Pheochromocytoma and Paraganglioma | PCPG | 82 |
| Prostate adenocarcinoma | PRAD | 351 |
| Sarcoma | SARC | 224 |
| Skin Cutaneous Melanoma | SKCM | 355 |
| Stomach adenocarcinoma | STAD | 392 |
| Testicular Germ Cell Tumors | TGCT | 122 |
| Thyroid carcinoma | THCA | 380 |
| Thymoma | THYM | 90 |
| Uterine Corpus Endometrial Carcinoma | UCEC | 439 |
| Uterine Carcinosarcoma | UCS | 48 |
| Uveal Melanoma | UVM | 12 |

**Supplementary Table S2.** Summary of the genes that the 12 pathways consist of.

| Pathway | Genes / Proteins |
|---|---|
| Apoptosis | BAD, BAK1, BAX, BCL2, BCL2L1, BID, BCL2L11, CASP7, BIRC2 |
| Breast reactive | CTNNB1, CAV1, GAPDH, MYH11, RAB11A, RAB11B, RBM15 |
| Cell cycle | CDK1, CCNB1, CCNE1, CCNE2, FOXM1, CDKN1B, PCNA |
| Core reactive | CTNNB1, CAV1, CLDN7, CDH1, RBM15 |
| DNA damage response | TP53BP1, ATM, BRCA2, CHEK1, CHEK2, XRCC5, MRE11A, TP53, RAD50, RAD51, XRCC1 |
| EMT | CTNNB1, CLDN7, COL6A1, CDH1, FN1, CDH2, SERPINE1 |
| Hormone receptor | AR, ESR1, PGR |
| Hormone signaling (Breast) | BCL2, GATA3, INPP4B |
| PI3K/AKT | AKT1, AKT2, AKT3, GSK3A, GSK3B, INPP4B, CDKN1B, AKT1S1, PTEN, TSC2 |
| RAS/MAPK | ARAF, JUN, RAF1, MAPK8, MAPK1, MAPK3, MAP2K1, MAPK14, RPS6KA1, YBX1 |
| RTK | EGFR, ERBB2, ERBB3, SHC1, SRC |
| TSC/mTOR | EIF4EBP1, MTOR, RPS6KB1, RB1, RPS6 |

**Supplementary Table S3.** Summary of cell line expression sample sizes according to lineages.

| Cancer Lineage | Number of Cell lines |
|---|---|
| bladder | 11 |
| blood | 101 |
| bone | 20 |
| brain | 6 |
| breast | 57 |
| colon | 35 |
| head and neck | 53 |
| kidney | 29 |
| liver | 17 |
| lung | 124 |
| ovary | 47 |
| pancreas | 20 |
| sarcoma | 29 |
| skin | 46 |
| stomach-oesophagus | 13 |
| uterus | 32 |

**Supplementary Table S4.** Summary of drug sensitivity data availability for cell lines.

| Cancer Lineage | Number of Cell lines with Drug Response Data | Average Number of Drug Responses Available / Cell line |
|---|---|---|
| bladder | 3 | 73 |
| blood | 4 | 132 |
| bone | 7 | 85 |
| brain | 1 | 88 |
| breast | 35 | 105 |
| colon | 23 | 96 |
| head and neck | 3 | 84 |
| kidney | 6 | 93 |
| liver | 13 | 75 |
| lung | 72 | 87 |
| ovary | 25 | 78 |
| pancreas | 17 | 82 |
| sarcoma | 4 | 64 |
| skin | 15 | 105 |
| stomach-oesophagus | 9 | 81 |
| uterus | 17 | 78 |

**Supplementary Table S5.** Major findings for pan-cancer, pathway-specific conserved and differential connectivity across lineages. Edges having EC > 50% of the total number of lineages for both patients and cell lines with a PPI score > 0.5 are marked in **bold**; the same with a PPI score < 0.5, which are marked in *italics*.

| Pathways | Edges with EC > 50% of Total Number of Lineages (Number of Lineages) |
|---|---|
| | **PPI Score > 0.5, EC > 50% among cell line lineages** |
| Apoptosis | BAX-BCL2(8), **BAX-BID(8)**, BCL2-BIRC2(9) |
| Breast reactive | CAV1-CTNNB1(11), GAPDH-MYH11(9) |
| Cell cycle | CCNE1-CDKN1B(8), CDK1-PCNA(9), **CDKN1B-PCNA(8)** |
| Core reactive | **CAV1-CTNNB1(8), CDH1-CLDN7(9)** |
| DNA damage response | ATM-TP53BP1(10), MRE11A-TP53BP1(8) |
| EMT | **CDH1-CLDN7(9), FN1-SERPINE1(8)** |
| Hormone receptor | **AR-ESR1(9), AR-PGR(10), ESR1-PGR(11)** |
| PI3K/AKT | AKT1S1-PTEN(8) |
| RAS/MAPK | ARAF-MAP2K1(8), JUN-RAF1(10) |
| RTK | **EGFR-ERBB2(9)**, ERBB2-SRC(9), SHC1-SRC(14) |
| TSC/mTOR | EIF4EBP1-MTOR(11), EIF4EBP1-RPS6(8), EIF4EBP1-RPS6KB1(8) |
| | **PPI Score < 0.5, EC > 50% among cell line lineages** |
| Breast reactive | MYH11-RBM15(9) |
| Cell cycle | *CCNE2-FOXM1(10)* |
| Core reactive | CLDN7-CTNNB1(9) |
| EMT | *CTNNB1-SERPINE1(8)* |
| Hormone signaling – Breast | BCL2-GATA3(9) |
| RAS/MAPK | ARAF-YBX1(9), MAPK14-RPS6KA1(8) |
| RTK | |
| TSC/mTOR | *RB1-RPS6(8)*, RB1-RPS6KB1(10) |
| | **PPI Score > 0.5, EC > 50% among patient lineages** |

| | |
|---|---|
| Apoptosis | BAD-BID(19), BAK1-BCL2L1(20), BAK1-BID(20), BAX-BCL2L1(16), **BAX-BID(17)**, BAX-CASP7(21), BCL2-BCL2L11(26), BCL2L1-BID(21), BCL2L11-BID(16) |
| Cell cycle | CCNB1-CCNE1(21), CCNB1-CDKN1B(19), CCNB1-FOXM1(24), CCNB1-PCNA(27), CCNE1-PCNA(18), CCNE2-CDK1(17), CCNE2-CDKN1B(22), CDK1-FOXM1(17), **CDKN1B-PCNA(16)** |
| Core reactive | **CAV1-CTNNB1(19), CDH1-CLDN7(25)**, CDH1-CTNNB1(27) |
| DNA damage response | ATM-RAD51(17), ATM-XRCC5(24), BRCA2-MRE11A(24), BRCA2-RAD51(17), BRCA2-TP53(17), CHEK1-CHEK2(20), CHEK1-MRE11A(20), CHEK1-TP53(16), CHEK2-MRE11A(27), CHEK2-TP53(21), MRE11A-RAD51(25), TP53-XRCC1(20), TP53BP1-XRCC5(28), XRCC1-XRCC5(17) |
| EMT | CDH1-CDH2(18), **CDH1-CLDN7(26)**, CDH1-CTNNB1(27), CDH2-CTNNB1(16), **FN1-SERPINE1(31)** |
| Hormone receptor | **AR-ESR1(19), AR-PGR(18), ESR1-PGR(26)** |
| PI3K/AKT | AKT1S1-TSC2(24) |
| RAS/MAPK | ARAF-JUN(16), JUN-MAP2K1(19), MAP2K1-RAF1(27), MAPK14-RAF1(17), RPS6KA1-YBX1(18) |
| RTK | **EGFR-ERBB2(22)**, EGFR-SHC1(27), EGFR-SRC(25), ERBB2-SHC1(16) |
| TSC/mTOR | MTOR-RPS6(28) |
| **PPI Score < 0.5, EC > 50% among patient lineages** | |
| Apoptosis | BID-CASP7(16) |
| Breast reactive | CAV1-MYH11(27), CAV1-RBM15(16), CTNNB1-RBM15(25) |
| Cell cycle | *CCNE2-FOXM1(17)* |
| EMT | CDH2-COL6A1(20), COL6A1-CTNNB1(20), COL6A1-FN1(19), COL6A1-SERPINE1(17), *CTNNB1-SERPINE1(17)* |
| PI3K/AKT | AKT1S1-CDKN1B(23) |
| RAS/MAPK | MAP2K1-RPS6KA1(18), MAP2K1-YBX1(21), RAF1-RPS6KA1(19) |
| RTK | |
| TSC/mTOR | EIF4EBP1-RB1(17), MTOR-RB1(22), *RB1-RPS6(20)* |

**Supplementary Table S6.** Connectivity scores (randomCS proportions) of the integrated cancer-specific networks. The connectivity scores with randomCS proportions < 0.15 are **bold**.

| Cancer Lineage | Apoptosis | Breast reactive | Cell cycle | Core reactive | DNA damage response | EMT | Hormone receptor | Hormone signaling (Breast) | PI3K/AKT | RAS/MAPK | RTK | TSC/mTOR |
|---|---|---|---|---|---|---|---|---|---|---|---|---|
| **Patient Cancers** | | | | | | | | | | | | |
| ACC | 0.28 (0.96) | 0.4 (0.689) | **0.67 (0.007)** | 0.4 (0.626) | 0.45 (0.164) | 0.5 (0.35) | 0.33 (0.872) | 0.67 (0.292) | 0.33 (0.813) | **0.57 (0.046)** | 0.67 (0.186) | 0.4 (0.594) |
| BLCA | 0.47 (0.837) | **0.77 (0.07)** | **0.71 (0.115)** | 0.7 (0.298) | 0.44 (0.864) | 0.57 (0.467) | 0.67 (0.757) | 0.67 (0.594) | 0.55 (0.589) | 0.52 (0.693) | 0.67 (0.567) | 0.35 (0.967) |
| BRCA | 0.58 (0.776) | **0.8 (0.099)** | 0.71 (0.372) | **0.9 (0.067)** | **0.69 (0.073)** | **0.79 (0.103)** | **1 (0)** | **1 (0)** | 0.67 (0.507) | 0.71 (0.163) | 1 (0) | **0.9 (0.099)** |
| CESC | 0.51 (0.23) | 0.5 (0.491) | 0.52 (0.325) | **0.75 (0.042)** | **0.55 (0.017)** | 0.52 (0.343) | **1 (0)** | **1 (0)** | 0.52 (0.31) | 0.45 (0.613) | 0.67 (0.341) | 0.3 (0.935) |
| CHOL | 0.53 (0.269) | 0.5 (0.55) | 0.45 (0.696) | **0.7 (0.148)** | **0.74 (0.001)** | 0.6 (0.238) | 0.5 (0.615) | 0.67 (0.455) | 0.33 (0.938) | 0.57 (0.187) | 0.67 (0.317) | 0.45 (0.653) |
| CORE | 0.58 (0.53) | 0.6 (0.592) | 0.52 (0.84) | 0.8 (0.178) | **0.63 (0.143)** | 0.69 (0.194) | **1 (0)** | **1 (0)** | 0.62 (0.483) | 0.59 (0.535) | 0.83 (0.296) | 0.7 (0.353) |
| DLBC | 0.4 (0.593) | 0.43 (0.488) | **0.52 (0.15)** | 0.5 (0.356) | **0.69 (0)** | 0.33 (0.805) | **1 (0)** | 0.5 (0.344) | **0.64 (0.022)** | 0.45 (0.353) | 0.42 (0.499) | 0.5 (0.335) |
| ESCA | 0.44 (0.494) | 0.57 (0.244) | **0.6 (0.117)** | **0.8 (0.034)** | 0.36 (0.875) | 0.5 (0.445) | **1 (0)** | **1 (0)** | **0.71 (0.006)** | **0.57 (0.113)** | 0.5 (0.637) | 0.4 (0.785) |
| GBM | 0.51 (0.521) | 0.6 (0.404) | 0.6 (0.361) | **0.8 (0.079)** | 0.54 (0.242) | 0.5 (0.634) | 0.67 (0.634) | 0.33 (0.944) | **0.67 (0.1)** | 0.5 (0.589) | 0.67 (0.447) | **0.75 (0.091)** |
| HNSC | 0.44 (0.596) | 0.47 (0.63) | 0.33 (0.936) | 0.6 (0.359) | **0.52 (0.094)** | 0.55 (0.261) | 0.67 (0.515) | 0.33 (0.89) | **0.67 (0.026)** | 0.48 (0.456) | 0.67 (0.281) | 0.5 (0.579) |
| KICH | 0.4 (0.897) | **0.7 (0.097)** | 0.62 (0.199) | 0.7 (0.255) | 0.48 (0.314) | 0.62 (0.175) | 0.67 (0.668) | 0 (1) | 0.55 (0.411) | 0.39 (0.934) | **1 (0)** | 0.55 (0.578) |
| KIRC | 0.54 (0.613) | 0.67 (0.303) | **0.81 (0.015)** | 0.6 (0.621) | 0.54 (0.563) | 0.5 (0.833) | **1 (0)** | 0.33 (0.959) | 0.64 (0.295) | **0.71 (0.04)** | 0.67 (0.577) | 0.7 (0.393) |
| KIRP | 0.56 (0.439) | 0.53 (0.745) | 0.64 (0.273) | 0.4 (0.955) | **0.58 (0.146)** | 0.55 (0.682) | 0.67 (0.738) | 0.67 (0.628) | 0.55 (0.657) | 0.62 (0.225) | 0.83 (0.268) | 0.6 (0.584) |
| LGG | 0.53 (0.902) | 0.73 (0.335) | 0.76 (0.162) | 0.3 (0.995) | 0.61 (0.476) | 0.55 (0.853) | 0.67 (0.782) | 0.33 (0.97) | 0.69 (0.35) | 0.62 (0.557) | 0.75 (0.39) | **0.9 (0.071)** |
| LIHC | 0.6 (0.172) | 0.33 (0.992) | 0.55 (0.62) | 0.5 (0.804) | 0.49 (0.577) | 0.62 (0.357) | 0.67 (0.782) | 0.67 (0.698) | 0.62 (0.332) | 0.5 (0.691) | 0.5 (0.886) | 0.6 (0.61) |
| LUAD | 0.58 (0.296) | 0.63 (0.348) | **0.74 (0.046)** | 0.65 (0.366) | 0.52 (0.421) | 0.64 (0.244) | 0.67 (0.681) | 0 (1) | 0.55 (0.523) | 0.45 (0.847) | 0.67 (0.529) | 0.7 (0.322) |
| LUSC | 0.46 (0.768) | 0.6 (0.411) | **0.67 (0.145)** | 0.7 (0.278) | 0.38 (0.954) | 0.57 (0.439) | **1 (0)** | 0.33 (0.946) | 0.64 (0.174) | **0.64 (0.111)** | 0.83 (0.205) | 0.5 (0.768) |
| MESO | 0.31 (0.913) | 0.5 (0.26) | 0.38 (0.648) | 0.5 (0.41) | 0.35 (0.689) | 0.48 (0.27) | **1 (0)** | 0.67 (0.308) | 0.38 (0.568) | 0.41 (0.421) | 0.58 (0.209) | 0.55 (0.216) |
| OV | 0.53 (0.217) | 0.5 (0.569) | 0.36 (0.937) | 0.6 (0.378) | 0.5 (0.238) | **0.62 (0.143)** | 0.67 (0.544) | **1 (0)** | 0.57 (0.246) | **0.7 (0.006)** | 0.67 (0.391) | 0.4 (0.878) |
| PAAD | 0.4 (0.795) | 0.57 (0.336) | 0.55 (0.303) | **0.8 (0.028)** | 0.45 (0.461) | 0.45 (0.663) | **1 (0)** | **1 (0)** | 0.57 (0.202) | 0.45 (0.634) | 0.67 (0.3) | 0.6 (0.342) |
| PCPG | 0.4 (0.693) | 0.57 (0.229) | 0.38 (0.832) | 0.5 (0.557) | 0.41 (0.517) | 0.55 (0.192) | 0.67 (0.487) | 0.33 (0.885) | 0.52 (0.243) | **0.64 (0.02)** | 0.33 (0.885) | 0.5 (0.568) |
| PRAD | 0.67 (0.192) | 0.73 (0.266) | 0.6 (0.721) | 0.65 (0.563) | 0.52 (0.816) | 0.57 (0.784) | 0.67 (0.756) | 0.67 (0.713) | 0.6 (0.664) | 0.59 (0.644) | 0.83 (0.352) | 0.6 (0.772) |
| SARC | 0.5 (0.367) | **0.77 (0.02)** | 0.48 (0.676) | 0.6 (0.395) | 0.43 (0.639) | 0.5 (0.534) | 0.67 (0.601) | 0.67 (0.489) | **0.69 (0.022)** | 0.52 (0.387) | 0.67 (0.396) | 0.4 (0.863) |
| SKCM | 0.39 (0.961) | 0.5 (0.731) | 0.62 (0.232) | 0.7 (0.244) | 0.51 (0.339) | **0.71 (0.062)** | 0.67 (0.682) | 0.33 (0.932) | 0.43 (0.884) | **0.66 (0.063)** | 0.5 (0.827) | **0.75 (0.135)** |
| STAD | 0.54 (0.692) | 0.53 (0.792) | 0.57 (0.673) | 0.7 (0.378) | 0.46 (0.934) | 0.69 (0.208) | **1 (0)** | 0.33 (0.959) | 0.43 (0.96) | 0.57 (0.53) | 0.83 (0.285) | 0.7 (0.373) |
| TGCT | 0.46 (0.596) | 0.6 (0.272) | 0.6 (0.218) | 0.55 (0.527) | **0.78 (0)** | 0.48 (0.602) | 0.33 (0.92) | 0.33 (0.901) | 0.55 (0.265) | 0.5 (0.442) | 0.17 (0.999) | 0.35 (0.928) |
| THCA | 0.53 (0.761) | 0.6 (0.606) | **0.74 (0.117)** | 0.75 (0.206) | 0.55 (0.611) | **0.79 (0.037)** | 0.67 (0.729) | 0.67 (0.657) | 0.57 (0.636) | **0.68 (0.15)** | 0.67 (0.57) | 0.7 (0.383) |
| THYM | 0.54 (0.153) | **0.63 (0.146)** | 0.43 (0.762) | 0.65 (0.174) | **0.67 (0)** | 0.55 (0.349) | 0.33 (0.932) | 0 (1) | 0.48 (0.542) | 0.46 (0.577) | 0.33 (0.928) | 0.5 (0.577) |
| UCEC | 0.44 (0.923) | 0.67 (0.256) | 0.64 (0.254) | **0.9 (0.023)** | 0.47 (0.808) | 0.67 (0.183) | **1 (0)** | 0 (1) | 0.62 (0.286) | **0.7 (0.053)** | 0.83 (0.181) | 0.45 (0.83) |
| UCS | 0.32 (0.797) | 0.4 (0.571) | 0.38 (0.576) | 0.6 (0.155) | 0.39 (0.337) | **0.52 (0.121)** | 0.33 (0.829) | 0.33 (0.803) | **0.62 (0.017)** | 0.48 (0.175) | 0.5 (0.465) | 0.3 (0.845) |
| UVM | 0.58 (0.596) | 0.37 (0.852) | 0.55 (0.506) | 0.3 (0.922) | 0.58 (0.906) | 0.52 (0.57) | 0 (1) | 0.67 (0.405) | 0.55 (0.507) | **0.77 (0.114)** | 0.67 (0.265) | 0.6 (0.319) |
| **Cell line Cancers** | | | | | | | | | | | | |
| bladder | 0.39 (0.767) | 0.63 (0.336) | 0.38 (0.867) | 0.8 (0.185) | 0.3 (0.897) | 0.57 (0.425) | 0.83 (0.786) | 0.83 (0.746) | 0.67 (0.242) | **0.71 (0.115)** | 0.75 (0.483) | 0.7 (0.321) |
| blood | 0.44 (0.71) | 0.5 (0.911) | 0.5 (0.774) | 0.7 (0.43) | 0.42 (0.495) | 0.6 (0.236) | 0.67 (0.999) | **1 (0)** | 0.55 (0.497) | 0.43 (0.927) | 0.83 (0.425) | 0.55 (0.926) |
| bone | **0.6 (0.038)** | 0.57 (0.427) | 0.4 (0.884) | 0.6 (0.507) | 0.35 (0.787) | 0.5 (0.546) | 0.67 (0.998) | **1 (0)** | 0.36 (0.969) | 0.38 (0.903) | 0.83 (0.202) | 0.6 (0.543) |
| brain | 0.33 (0.168) | 0.7 (0.2) | 0.55 (0.19) | 0.65 (0.753) | 0.13 (0.584) | 0.4 (0.701) | **1 (0)** | **1 (0)** | 0.52 (0.233) | 0.41 (0.2) | 0.5 (0.956) | 0.65 (0.723) |
| breast | 0.47 (0.398) | 0.43 (0.973) | 0.52 (0.497) | 0.6 (0.678) | 0.45 (0.348) | 0.4 (0.961) | 0.67 (0.991) | 0.67 (0.982) | 0.45 (0.844) | 0.46 (0.648) | 0.58 (0.85) | 0.6 (0.649) |
| colon | 0.49 (0.367) | 0.5 (0.546) | 0.31 (0.976) | **0.8 (0.04)** | 0.48 (0.27) | 0.43 (0.747) | **1 (0)** | **1 (0)** | 0.38 (0.848) | **0.59 (0.101)** | 0.67 (0.327) | 0.5 (0.544) |
| head and neck | 0.42 (0.675) | 0.53 (0.532) | 0.5 (0.492) | 0.65 (0.323) | 0.4 (0.501) | 0.4 (0.917) | 0.67 (0.971) | **1 (0)** | 0.38 (0.97) | 0.38 (0.908) | 0.5 (0.976) | 0.65 (0.381) |
| kidney | 0.51 (0.222) | 0.4 (0.978) | 0.57 (0.224) | 0.45 (0.968) | 0.35 (0.881) | 0.38 (0.971) | 0.83 (0.521) | 0.83 (0.525) | 0.55 (0.354) | **0.61 (0.063)** | 0.83 (0.208) | 0.55 (0.794) |
| liver | 0.38 (0.904) | 0.33 (0.995) | 0.43 (0.885) | 0.5 (0.835) | 0.52 (0.239) | **0.71 (0.045)** | 0.83 (0.569) | **1 (0)** | **0.64 (0.13)** | 0.52 (0.355) | 0.58 (0.761) | 0.75 (0.164) |
| lung | 0.51 (0.335) | 0.57 (0.576) | **0.62 (0.15)** | 0.65 (0.474) | 0.46 (0.44) | 0.55 (0.501) | **1 (0)** | 0.67 (0.969) | 0.48 (0.871) | 0.55 (0.297) | 0.83 (0.281) | 0.65 (0.449) |
| ovary | 0.47 (0.633) | 0.53 (0.661) | 0.62 (0.153) | 0.55 (0.79) | 0.46 (0.532) | 0.52 (0.597) | 0.67 (0.985) | 0.83 (0.389) | 0.55 (0.487) | 0.34 (0.998) | 0.58 (0.815) | 0.5 (0.926) |
| pancreas | 0.35 (0.894) | 0.4 (0.746) | 0.36 (0.834) | 0.6 (0.232) | **0.55 (0.141)** | 0.36 (0.831) | 0.67 (0.521) | 0.67 (0.396) | 0.55 (0.191) | **0.55 (0.149)** | 0.17 (0.992) | 0.6 (0.206) |
| sarcoma | 0.49 (0.477) | 0.63 (0.216) | 0.55 (0.375) | 0.5 (0.895) | 0.44 (0.653) | 0.4 (0.921) | 0.83 (0.439) | 0.67 (0.962) | 0.52 (0.5) | 0.59 (0.154) | 0.67 (0.67) | 0.6 (0.577) |
| skin | 0.4 (0.826) | 0.5 (0.806) | 0.52 (0.475) | 0.6 (0.713) | 0.35 (0.904) | **0.62 (0.123)** | 0.67 (0.995) | **1 (0)** | 0.57 (0.253) | 0.52 (0.303) | 0.67 (0.777) | 0.6 (0.668) |
| stomach-oesophagus | 0.53 (0.856) | 0.57 (0.755) | 0.62 (0.585) | 0.65 (0.59) | 0.63 (0.579) | 0.62 (0.609) | 0.83 (0.447) | 0 (1) | 0.64 (0.53) | 0.46 (0.948) | **1 (0)** | 0.55 (0.81) |
| uterus | 0.5 (0.338) | 0.57 (0.402) | 0.45 (0.741) | 0.65 (0.274) | 0.42 (0.695) | **0.62 (0.146)** | 0.67 (0.953) | 0.83 (0.346) | 0.6 (0.163) | 0.41 (0.823) | 0.5 (0.954) | 0.6 (0.491) |

**Supplementary Table S7.** Percentages of sample-sample pairs for each patient cancer (in rows) by cell line cancer (in columns) combination that have a Pearson's correlation of modulus 0.9 or higher between their network aberration score vectors across the 12 pathways.

| | bladder | blood | bone | brain | breast | colon | head and neck | kidney | liver | lung | ovary | pancreas | sarcoma | skin | stomach-oesophagus | uterus |
|---|---|---|---|---|---|---|---|---|---|---|---|---|---|---|---|---|
| **ACC** | 0 | 29.51 | 0 | 19.2 | 20.94 | 7.33 | 16.49 | 4.35 | 58.57 | 2.88 | 0.51 | 77.5 | 7.12 | 0.99 | 18.73 | 6.52 |
| **BLCA** | 0 | 0.13 | 0 | 0 | 26.12 | 0 | 17.83 | 32.06 | 4.1 | 28.57 | 13.07 | 0.77 | 11.6 | 28.64 | 0 | 0.56 |
| **BRCA** | 0 | 66.94 | 7.1 | 53.08 | 94.51 | 1.31 | 91.98 | 83.18 | 88.43 | 86.92 | 54.7 | 96.94 | 94.14 | 83.21 | 46.89 | 62.5 |
| **CESC** | 0 | 75.2 | 0.26 | 71.83 | 87.13 | 15.44 | 76.46 | 51 | 69.21 | 64.2 | 37.35 | 79.77 | 76.89 | 53.19 | 63.38 | 71.24 |
| **CHOL** | 0 | 0 | 0 | 0 | 0 | 0 | 0 | 0 | 0 | 0 | 0 | 0 | 0 | 0 | 0 | 0 |
| **CORE** | 0 | 74.15 | 22.81 | 63.31 | 82.38 | 1.31 | 88.33 | 66.54 | 75.42 | 84.22 | 48.31 | 83.27 | 89.61 | 72.24 | 61.51 | 51.71 |
| **DLBC** | 0 | 0 | 0 | 0 | 0 | 0 | 0 | 0 | 0 | 0 | 0 | 0 | 0 | 0 | 0 | 0 |
| **ESCA** | 0 | 42.05 | 0 | 47.88 | 31.86 | 9.41 | 20.25 | 87.99 | 17.65 | 36.41 | 3.24 | 62.66 | 64.45 | 42.67 | 2.44 | 42.61 |
| **GBM** | 0 | 68.37 | 0.04 | 77.3 | 86.4 | 7.88 | 81.32 | 36.22 | 64.78 | 65.26 | 65.16 | 93.06 | 65.7 | 57.27 | 55.04 | 43.47 |
| **HNSC** | 0 | 65 | 0.17 | 49.75 | 41.02 | 50.19 | 60.36 | 23.48 | 76.3 | 14.03 | 20.68 | 99.24 | 61.17 | 27.83 | 51.12 | 29.99 |
| **KICH** | 0 | 52.27 | 0 | 72.49 | 8.05 | 63.45 | 11.65 | 0.44 | 11.67 | 3.43 | 4.42 | 60.16 | 33.55 | 2.69 | 34.31 | 19.79 |
| **KIRC** | 0 | 78.89 | 0 | 74.48 | 45.72 | 43.55 | 35.81 | 79.05 | 19.15 | 55.14 | 5.46 | 90.9 | 84.6 | 49.08 | 7.73 | 42.48 |
| **KIRP** | 0 | 43.25 | 6.96 | 73.58 | 49.12 | 25.39 | 53.46 | 44.64 | 39.44 | 44.39 | 5.78 | 72.24 | 59 | 21.11 | 54.13 | 44.69 |
| **LGG** | 0 | 89.32 | 0.02 | 86.19 | 81.06 | 15.99 | 85.18 | 46.81 | 76.03 | 67.62 | 55.8 | 97.48 | 80.24 | 55.63 | 50 | 43.63 |
| **LIHC** | 0 | 99.81 | 28.23 | 17.12 | 38.07 | 4.75 | 31.44 | 86.77 | 62.66 | 85.04 | 20.33 | 67.07 | 94.36 | 57.46 | 24.21 | 88.04 |
| **LUAD** | 0 | 48.76 | 0 | 62.34 | 48.78 | 5.97 | 46.58 | 31.22 | 33.38 | 51.58 | 20.31 | 40.43 | 44.1 | 28.4 | 34.81 | 34.65 |
| **LUSC** | 0 | 31.11 | 0.12 | 18.05 | 31.04 | 2.07 | 19.12 | 49.31 | 20.27 | 33.76 | 12.8 | 27.2 | 35.61 | 32.68 | 2.58 | 27.3 |
| **MESO** | 0 | 11 | 0 | 73.77 | 0.14 | 21.83 | 1.73 | 0.11 | 13.89 | 0 | 0.03 | 25.25 | 16.11 | 0.11 | 26.61 | 8.81 |
| **OV** | 0 | 32.74 | 43.03 | 37.86 | 51.81 | 1.05 | 67.22 | 89.4 | 58.59 | 63.39 | 12.38 | 82.65 | 76.61 | 60.7 | 19.36 | 35.89 |
| **PAAD** | 0 | 53.17 | 0.9 | 25.41 | 95.21 | 0.75 | 91.88 | 58.85 | 73.92 | 90.21 | 77.01 | 61.31 | 51.64 | 73.24 | 58.58 | 57.84 |
| **PCPG** | 0 | 34.3 | 5.3 | 43.29 | 20.77 | 27.8 | 18.57 | 76.45 | 22.96 | 17.76 | 0.6 | 83.05 | 80.36 | 34.09 | 9.66 | 45.05 |
| **PRAD** | 0 | 13.62 | 7.41 | 53.37 | 84.51 | 1.39 | 92.31 | 50.64 | 40.02 | 75.45 | 21.13 | 69.79 | 63.27 | 50.86 | 59.68 | 30.93 |
| **SARC** | 0 | 59.95 | 0.42 | 62.2 | 37.82 | 66.38 | 30.26 | 21.37 | 7.17 | 6.14 | 4.36 | 80.2 | 88.45 | 28.78 | 62.02 | 79.63 |
| **SKCM** | 0 | 47.21 | 0.03 | 39.44 | 3.38 | 15.94 | 3.27 | 18 | 36.44 | 11.93 | 1.58 | 34.35 | 38.79 | 6.64 | 7.89 | 26.4 |
| **STAD** | 0 | 9.29 | 13.04 | 26.53 | 61.6 | 0.05 | 78.34 | 71.24 | 52.49 | 83.63 | 13.36 | 34.27 | 55.93 | 48.78 | 9.14 | 6.78 |
| **TGCT** | 0 | 0 | 0 | 0 | 0 | 0 | 0 | 0 | 0 | 0 | 0 | 0 | 0 | 0 | 0 | 0 |
| **THCA** | 0 | 73.08 | 0.63 | 62.06 | 48.93 | 4.32 | 39.88 | 48.76 | 92.65 | 52.74 | 10.62 | 89.68 | 52.15 | 25.85 | 14.31 | 24.11 |
| **THYM** | 0 | 0 | 0 | 0 | 0 | 0 | 0 | 0 | 0 | 0 | 0 | 0 | 0 | 0 | 0 | 0 |
| **UCEC** | 0 | 73.34 | 0.63 | 40.77 | 40.52 | 9.39 | 21.44 | 79.24 | 53.75 | 50.12 | 22.53 | 70.42 | 77.27 | 57.54 | 7.24 | 58.52 |
| **UCS** | 0 | 49.38 | 0 | 4.17 | 0.91 | 31.96 | 0.31 | 16.74 | 41.18 | 0.13 | 0.18 | 84.38 | 21.19 | 2.49 | 10.42 | 32.88 |
| **UVM** | 0 | 17.66 | 0 | 4.17 | 5.7 | 14.52 | 6.76 | 34.2 | 65.2 | 2.49 | 0 | 98.75 | 7.18 | 0.54 | 5.13 | 8.07 |
| **UVM** | 0 | 7.01 | 0 | 29.17 | 2.19 | 2.86 | 7.08 | 1.15 | 46.08 | 1.14 | 0 | 96.25 | 1.44 | 2.72 | 10.9 | 13.8 |

**Supplementary Table S8.** Membership of samples by cancer lineages in the 29 clusters obtained from hierarchical clustering on the network aberration scores matrix. The color codes are as follows – patient cancers, cell line cancers, clusters with notable membership from both the model systems.

| Cancer Lineages | 1 | 2 | 3 | 4 | 5 | 6 | 7 | 8 | 9 | 10 | 11 | 12 | 13 | 14 | 15 | 16 | 17 | 18 | 19 | 20 | 21 | 22 | 23 | 24 | 25 | 26 | 27 | 28 | 29 |
|---|---|---|---|---|---|---|---|---|---|---|---|---|---|---|---|---|---|---|---|---|---|---|---|---|---|---|---|---|---|
| ACC | 30 | 15 | 1 | 0 | 0 | 0 | 0 | 0 | 0 | 0 | 0 | 0 | 0 | 0 | 0 | 0 | 0 | 0 | 0 | 0 | 0 | 0 | 0 | 0 | 0 | 0 | 0 | 0 | 0 |
| BLCA | 0 | 0 | 0 | 76 | 50 | 209 | 8 | 0 | 0 | 0 | 0 | 0 | 0 | 0 | 0 | 0 | 0 | 0 | 0 | 0 | 0 | 0 | 0 | 0 | 0 | 0 | 0 | 0 | 0 |
| BRCA | 0 | 0 | 0 | 0 | 0 | 0 | 0 | 455 | 57 | 230 | 114 | 5 | 17 | 0 | 0 | 0 | 0 | 0 | 0 | 0 | 0 | 0 | 0 | 0 | 0 | 0 | 0 | 0 | 0 |
| CESC | 87 | 0 | 0 | 0 | 0 | 0 | 0 | 0 | 0 | 0 | 0 | 0 | 0 | 1 | 82 | 1 | 0 | 0 | 0 | 0 | 0 | 0 | 0 | 0 | 0 | 0 | 0 | 0 | 0 |
| CHOL | 0 | 0 | 0 | 0 | 0 | 0 | 30 | 0 | 0 | 0 | 0 | 0 | 0 | 0 | 0 | 0 | 0 | 0 | 0 | 0 | 0 | 0 | 0 | 0 | 0 | 0 | 0 | 0 | 0 |
| CORE | 0 | 0 | 0 | 17 | 0 | 0 | 0 | 1 | 47 | 298 | 27 | 1 | 54 | 14 | 0 | 0 | 32 | 0 | 0 | 0 | 0 | 0 | 0 | 0 | 0 | 0 | 0 | 0 | 0 |
| DLBC | 0 | 0 | 0 | 0 | 0 | 0 | 6 | 0 | 0 | 0 | 0 | 0 | 0 | 0 | 0 | 0 | 0 | 27 | 0 | 0 | 0 | 0 | 0 | 0 | 0 | 0 | 0 | 0 | 0 |
| ESCA | 24 | 1 | 0 | 0 | 21 | 0 | 0 | 0 | 0 | 0 | 0 | 0 | 0 | 0 | 55 | 21 | 0 | 0 | 4 | 0 | 0 | 0 | 0 | 0 | 0 | 0 | 0 | 0 | 0 |
| GBM | 12 | 0 | 0 | 9 | 0 | 0 | 1 | 0 | 1 | 0 | 0 | 4 | 0 | 0 | 45 | 2 | 0 | 0 | 0 | 154 | 4 | 0 | 0 | 0 | 0 | 0 | 0 | 0 | 0 |
| HNSC | 0 | 0 | 0 | 78 | 0 | 0 | 0 | 0 | 0 | 0 | 0 | 0 | 0 | 0 | 122 | 0 | 0 | 0 | 0 | 3 | 0 | 0 | 0 | 0 | 0 | 0 | 0 | 0 | 0 |
| KICH | 3 | 0 | 0 | 0 | 0 | 0 | 0 | 0 | 0 | 0 | 0 | 0 | 0 | 0 | 5 | 2 | 0 | 5 | 0 | 0 | 0 | 5 | 43 | 0 | 0 | 0 | 0 | 0 | 0 |
| KIRC | 1 | 0 | 0 | 0 | 0 | 0 | 0 | 0 | 6 | 0 | 0 | 0 | 0 | 0 | 3 | 78 | 0 | 0 | 0 | 2 | 125 | 56 | 0 | 87 | 87 | 24 | 0 | 0 | 0 |
| KIRP | 7 | 0 | 0 | 0 | 0 | 0 | 0 | 0 | 0 | 0 | 0 | 82 | 0 | 0 | 41 | 12 | 0 | 0 | 0 | 15 | 35 | 1 | 0 | 20 | 4 | 0 | 0 | 0 | 0 |
| LGG | 2 | 0 | 0 | 0 | 4 | 0 | 0 | 0 | 1 | 0 | 0 | 5 | 0 | 0 | 2 | 7 | 0 | 0 | 0 | 44 | 61 | 23 | 0 | 283 | 0 | 0 | 0 | 0 | 0 |
| LIHC | 0 | 0 | 0 | 0 | 0 | 0 | 0 | 0 | 0 | 0 | 0 | 0 | 0 | 0 | 0 | 0 | 0 | 0 | 150 | 0 | 0 | 0 | 0 | 0 | 0 | 0 | 33 | 1 | 0 |
| LUAD | 47 | 0 | 0 | 0 | 0 | 75 | 0 | 0 | 1 | 0 | 0 | 28 | 0 | 0 | 13 | 33 | 0 | 2 | 0 | 27 | 0 | 31 | 0 | 54 | 0 | 51 | 0 | 0 | 0 |
| LUSC | 0 | 0 | 0 | 0 | 16 | 19 | 0 | 0 | 0 | 0 | 0 | 5 | 0 | 0 | 0 | 39 | 0 | 13 | 0 | 0 | 0 | 53 | 29 | 0 | 50 | 101 | 0 | 0 | 0 |
| MESO | 7 | 54 | 0 | 0 | 0 | 0 | 0 | 0 | 0 | 0 | 0 | 0 | 0 | 0 | 0 | 0 | 0 | 0 | 0 | 0 | 0 | 0 | 0 | 0 | 0 | 0 | 0 | 0 | 0 |
| OV | 0 | 0 | 0 | 49 | 0 | 0 | 0 | 0 | 2 | 61 | 87 | 19 | 155 | 40 | 2 | 0 | 16 | 0 | 0 | 0 | 0 | 0 | 0 | 0 | 0 | 0 | 0 | 0 | 0 |
| PAAD | 0 | 0 | 1 | 97 | 0 | 0 | 24 | 0 | 0 | 0 | 0 | 0 | 0 | 0 | 0 | 0 | 0 | 0 | 0 | 0 | 0 | 0 | 0 | 0 | 0 | 0 | 0 | 0 | 0 |
| PCPG | 26 | 56 | 0 | 0 | 0 | 0 | 0 | 0 | 0 | 0 | 0 | 0 | 0 | 0 | 0 | 0 | 0 | 0 | 0 | 0 | 0 | 0 | 0 | 0 | 0 | 0 | 0 | 0 | 0 |
| PRAD | 0 | 0 | 0 | 0 | 1 | 0 | 0 | 0 | 53 | 84 | 90 | 55 | 10 | 13 | 0 | 0 | 0 | 0 | 0 | 1 | 0 | 0 | 0 | 44 | 0 | 0 | 0 | 0 | 0 |
| SARC | 0 | 0 | 0 | 0 | 0 | 0 | 0 | 0 | 0 | 0 | 0 | 0 | 0 | 0 | 67 | 157 | 0 | 0 | 0 | 0 | 0 | 0 | 0 | 0 | 0 | 0 | 0 | 0 | 0 |
| SKCM | 25 | 0 | 0 | 0 | 0 | 29 | 0 | 0 | 0 | 0 | 0 | 0 | 0 | 0 | 5 | 6 | 0 | 12 | 0 | 0 | 5 | 125 | 42 | 0 | 0 | 106 | 0 | 0 | 0 |
| STAD | 0 | 0 | 0 | 0 | 224 | 0 | 0 | 0 | 0 | 0 | 0 | 87 | 0 | 75 | 0 | 0 | 0 | 0 | 0 | 6 | 0 | 0 | 0 | 0 | 0 | 0 | 0 | 0 | 0 |
| TGCT | 0 | 0 | 0 | 0 | 0 | 28 | 0 | 0 | 0 | 0 | 0 | 0 | 0 | 0 | 0 | 0 | 0 | 87 | 0 | 0 | 0 | 0 | 0 | 0 | 0 | 7 | 0 | 0 | 0 |
| THCA | 0 | 0 | 0 | 0 | 9 | 0 | 0 | 0 | 0 | 0 | 0 | 3 | 0 | 0 | 6 | 1 | 0 | 0 | 0 | 3 | 31 | 72 | 0 | 230 | 25 | 0 | 0 | 0 | 0 |
| THYM | 0 | 0 | 0 | 0 | 0 | 0 | 0 | 0 | 0 | 0 | 0 | 0 | 0 | 0 | 0 | 0 | 0 | 90 | 0 | 0 | 0 | 0 | 0 | 0 | 0 | 0 | 0 | 0 | 0 |
| UCEC | 26 | 0 | 0 | 0 | 0 | 6 | 0 | 0 | 0 | 0 | 0 | 0 | 0 | 0 | 2 | 49 | 0 | 0 | 0 | 78 | 20 | 105 | 15 | 0 | 108 | 30 | 0 | 0 | 0 |
| UCS | 0 | 48 | 0 | 0 | 0 | 0 | 0 | 0 | 0 | 0 | 0 | 0 | 0 | 0 | 0 | 0 | 0 | 0 | 0 | 0 | 0 | 0 | 0 | 0 | 0 | 0 | 0 | 0 | 0 |
| UVM | 0 | 10 | 0 | 0 | 0 | 0 | 0 | 0 | 0 | 0 | 0 | 0 | 0 | 0 | 0 | 0 | 0 | 0 | 0 | 0 | 0 | 0 | 0 | 0 | 0 | 0 | 0 | 2 | 0 |
| bladder | 0 | 0 | 11 | 0 | 0 | 0 | 0 | 0 | 0 | 0 | 0 | 0 | 0 | 0 | 0 | 0 | 0 | 0 | 0 | 0 | 0 | 0 | 0 | 0 | 0 | 0 | 0 | 0 | 0 |
| blood | 0 | 0 | 17 | 0 | 0 | 0 | 0 | 0 | 0 | 0 | 0 | 0 | 0 | 0 | 0 | 0 | 0 | 0 | 36 | 0 | 0 | 0 | 0 | 0 | 0 | 0 | 0 | 48 | 0 |
| bone | 0 | 0 | 0 | 0 | 0 | 0 | 0 | 0 | 0 | 0 | 0 | 0 | 0 | 0 | 0 | 0 | 0 | 0 | 0 | 0 | 0 | 0 | 0 | 0 | 0 | 0 | 0 | 15 | 5 |
| brain | 0 | 0 | 6 | 0 | 0 | 0 | 0 | 0 | 0 | 0 | 0 | 0 | 0 | 0 | 0 | 0 | 0 | 0 | 0 | 0 | 0 | 0 | 0 | 0 | 0 | 0 | 0 | 0 | 0 |
| breast | 0 | 0 | 7 | 19 | 0 | 0 | 0 | 0 | 0 | 0 | 0 | 0 | 0 | 0 | 0 | 0 | 0 | 0 | 0 | 0 | 0 | 0 | 0 | 0 | 0 | 0 | 0 | 25 | 6 |
| colon | 0 | 9 | 0 | 0 | 0 | 0 | 0 | 0 | 0 | 0 | 0 | 0 | 0 | 0 | 0 | 0 | 0 | 0 | 0 | 0 | 0 | 0 | 25 | 0 | 0 | 0 | 0 | 0 | 1 |
| head and neck | 0 | 0 | 4 | 38 | 0 | 0 | 0 | 0 | 0 | 0 | 0 | 0 | 0 | 1 | 0 | 0 | 0 | 0 | 0 | 0 | 0 | 0 | 0 | 0 | 0 | 0 | 0 | 6 | 4 |
| kidney | 0 | 0 | 4 | 10 | 0 | 0 | 0 | 0 | 0 | 0 | 0 | 0 | 0 | 0 | 0 | 0 | 0 | 0 | 0 | 0 | 0 | 0 | 0 | 0 | 0 | 0 | 0 | 11 | 4 |
| liver | 0 | 0 | 2 | 0 | 0 | 0 | 0 | 0 | 0 | 0 | 0 | 0 | 0 | 0 | 0 | 0 | 0 | 0 | 0 | 0 | 0 | 0 | 0 | 0 | 0 | 0 | 0 | 13 | 2 |
| lung | 0 | 0 | 7 | 44 | 0 | 0 | 0 | 0 | 50 | 0 | 0 | 0 | 0 | 0 | 0 | 0 | 0 | 0 | 0 | 0 | 0 | 0 | 0 | 0 | 0 | 0 | 0 | 14 | 9 |
| ovary | 0 | 0 | 2 | 38 | 0 | 0 | 0 | 0 | 0 | 0 | 0 | 0 | 0 | 0 | 0 | 0 | 0 | 0 | 0 | 0 | 0 | 0 | 0 | 0 | 0 | 0 | 0 | 3 | 4 |
| pancreas | 0 | 14 | 1 | 4 | 0 | 0 | 0 | 0 | 0 | 0 | 0 | 0 | 0 | 0 | 0 | 0 | 0 | 0 | 0 | 0 | 0 | 0 | 0 | 0 | 0 | 0 | 0 | 0 | 1 |
| sarcoma | 0 | 0 | 12 | 0 | 0 | 0 | 0 | 0 | 0 | 0 | 0 | 0 | 0 | 0 | 0 | 0 | 0 | 0 | 10 | 0 | 0 | 0 | 0 | 0 | 0 | 0 | 0 | 7 | 0 |
| skin | 0 | 0 | 7 | 0 | 0 | 0 | 0 | 0 | 0 | 0 | 0 | 0 | 0 | 0 | 0 | 0 | 0 | 0 | 0 | 0 | 0 | 0 | 0 | 0 | 0 | 0 | 0 | 11 | 6 |
| stomach-oesophagus | 0 | 0 | 4 | 0 | 0 | 0 | 0 | 0 | 0 | 0 | 0 | 0 | 0 | 0 | 5 | 0 | 0 | 0 | 0 | 0 | 0 | 0 | 0 | 0 | 0 | 0 | 0 | 3 | 1 |
| uterus | 0 | 0 | 0 | 0 | 0 | 0 | 0 | 0 | 0 | 0 | 0 | 0 | 0 | 0 | 0 | 0 | 0 | 0 | 20 | 0 | 0 | 0 | 0 | 0 | 0 | 0 | 0 | 7 | 5 |

**Supplementary Table S9.** Shared memberships of cell lines and cancer patients in clusters obtained from hierarchical clustering on the network aberration scores, along with the pathways driving the shared presence.

| Cluster | Cell Line Lineages | Patient Tumors | Driving Pathways |
|---|---|---|---|
| C2 | pancreas (70%)<br>colon (26%) | Mesothelioma (89%)<br>Uveal Melanoma (83%)<br>Pheochromocytoma and Paraganglioma (68%)<br>Adrenocortical carcinoma (33%) | RAS/MAPK |
| C4 | ovary (81%)<br>head and neck (72%)<br>skin (48%)<br>lung (35%)<br>kidney (34%)<br>breast (33%)<br>pancreas (20%) | Pancreatic adenocarcinoma (80%)<br>Head and Neck squamous cell carcinoma (38%)<br>Bladder Urothelial Carcinoma (22%)<br>Ovarian serous cystadenocarcinoma (11%) | Apoptosis<br>DNA Damage Response |
| C9 | lung (40%) | Prostate adenocarcinoma (15%) | Apoptosis<br>Breast Reactive<br>Core Reactive<br>DNA Damage Response |
| C15 | stomach-oesophagus (38%) | Head and Neck squamous cell carcinoma (60%)<br>Cervical squamous cell carcinoma and endocervical adenocarcinoma (48%)<br>Esophageal carcinoma (40%)<br>Sarcoma (30%)<br>Kidney renal papillary cell carcinoma (19%)<br>Glioblastoma multiforme (19%) | DNA Damage Response<br>PI3K/AKT |
| C19 | uterus (62%)<br>blood (36%)<br>sarcoma (34%) | Liver hepatocellular carcinoma (82%) | DNA Damage Response<br>PI3K/AKT<br>RAS/MAPK |
| C23 | colon (71%) | Kidney Chromophobe (68%) | Apoptosis<br>Cell Cycle<br>EMT |

**Supplementary Table S10.** Summary of Bayesian additive regression tree models for each cancer type and each drug. Models are fit using cell lines' drug sensitivity and network aberration scores. Then the patient drug responses for a cancer type are predicted using the model for that drug and the cell line cancer lineage for the same tissue. The response rate is defined as the proportion of samples for that cancer type with a >0.5 predicted response with respect to that drug. Drugs marked with an asterisk (*) have been or are being investigated through clinical trials on cancers of the corresponding tissues (see Discussion section of the paper for references).

| Patient Cancer | Drugs/Compounds (Response Rates) | Top 3 Predictive Pathways in Cell Line Training Models | Targets |
|---|---|---|---|
| **BRCA** | Ibrutinib* (100%) | RAS/MAPK, PI3K/AKT, EMT | BTK |
| **CORE** | Lapatinib* (100%) | RAS/MAPK, Cell cycle, EMT | ERBB2, EGFR |
| | KU.0060648 (98.2%) | Apoptosis, Hormone signaling, RAS/MAPK | PI3K, DNA-PK |
| | PD318088 (96.7%) | RTK, PI3K/AKT, Core reactive | MEK1/2 |
| | Canertinib (94.9%) | Core reactive, RAS/MAPK, Hormone signaling | EGFR, HER-2, ErbB-4 |
| | Neratinib (93.5%) | Cell cycle, RTK, EMT | Her2, EGFR |
| | Afatinib (93.5%) | DNA damage response, Hormone signaling, Breast reactive | ERBB2, EGFR |
| | Navitoclax (68.4%) | EMT, Breast reactive, Cell cycle | BCL2, BCL-XL, BCL-W |
| **LIHC** | PD318088 (100%) | PI3K/AKT, Cell cycle, EMT | MEK1/2 |
| | UNC0638 (100%) | Apoptosis, PI3K/AKT, DNA damage response | G9a, GLP |
| | Linifanib (100%) | PI3K/AKT, EMT, RAS/MAPK | VEGFR1-3, CSF1R, FLT3, KIT |
| | Pandacostat (99.5%) | Breast reactive, RTK, RAS/MAPK | HDAC1-9 |
| **PAAD** | NPC.26 (100%) | Cell cycle, Breast reactive, TSC/mTOR | mPTP |
| | Trametinib (72.1%) | Apoptosis, TSC/mTOR, Cell cycle | MEK1/2 |

| | | | |
|---|---|---|---|
| | BRD9876 (62.3%) | RAS/MAPK, TSC/mTOR, PI3K/AKT | Eg5 |
| | UNC0638 (58.2%) | EMT, Breast reactive, Cell cycle | G9a and GLP methyltransferases |
| | Methotrexate (50.8%) | Apoptosis, Cell Cycle, Breast reactive | Antimetabolite |
| **SKCM** | PLX.4032 (100%) | EMT, DNA damage response, PI3K/AKT | B-Raf |
| | GDC.0879 (100%) | PI3K/AKT, EMT, Cell cycle | B-Raf |
| | PLX.4720 (81.4%) | RAS/MAPK, Apoptosis, PI3K/AKT | B-Raf |
| | VAF.347 (52.7%) | PI3K/AKT, DNA damage response, RAS/MAPK | AhR |
| **UCEC** | BRD.K97651142 (74.9%) | Breast reactive, RAS/MAPK, RTK | EGFR, ERBB2 |
